# The orbitofrontal cortex updates beliefs for state inference

**DOI:** 10.1101/2024.10.29.620879

**Authors:** Shannon S. Schiereck, Danilo Trinidad Pérez-Rivera, Andrew Mah, Margaret L. DeMaegd, David Hocker, Royall McMahon Ward, Cristina Savin, Christine M. Constantinople

## Abstract

While the orbitofrontal cortex (OFC) is implicated in learning and inferring latent states, the precise computation performed by OFC for state inference is unclear. Here we show that rat OFC updates beliefs about states, and this process is decipherable from OFC dynamics in rats performing state inference, but not alternative strategies. We trained rats to perform a temporal wagering task with hidden reward states. Well-trained rats used state inference when deciding how long to wait for rewards, and OFC inactivations impaired belief updating about states. Electrophysiology and novel population analysis methods identified latent neural factors reflecting inferred states in rats performing inference, but not other strategies. Neural firing rates and latent population factors showed abrupt changes following trials that were informative of state transitions. These results identify a precise computation performed by OFC, and reveal neural signatures of inference.

## Introduction

To survive in dynamic environments, animals cannot exclusively rely on learned stimulus-response associations, but must generalize and form inferences about the world; this process is among the most important and interesting cognitive operations that nervous systems perform. The orbitofrontal cortex (OFC) in rodents and primates is implicated in state inference when task contingencies are partially observable^1–6^, when values must be inferred based on high-order associations^7^, and as latent states are learned from experience^8^. How local circuit dynamics in OFC support state inference, however, remains unclear.

A key ingredient of statistical (Bayesian) inference is computing and combining belief distributions, that is, probability distributions over observations and hypothesized possible states. Behaviorally, humans and animals have been shown to combine prior beliefs with sensory evidence in perceptual and sensorimotor domains^9,10^. Neurally, progress has been made in characterizing neural representations of beliefs over sensory stimuli^11^ or perceptual judgments^12–16^ in psychophysical tasks. An outstanding question is whether or how neural representations of subjective beliefs might extend to more cognitive scenarios, beyond perceptual inference of physical sensory stimuli that exist in the world. How neural circuits compute and represent beliefs over abstract, latent states of the environment, is unresolved.

Most cognitive functions, including state inference, are compositional, in that they reflect multiple sub-computations that in principle could be supported by distinct neural circuits. For instance, state inference requires representing prior beliefs, processing sensory observations from the environment, combining these observations with prior beliefs, recursively updating prior beliefs, and working memory depending on the timescale of the task. It is currently unclear which precise sub-computation is supported by OFC as animals learn and infer latent states. This is a hard problem, in part because behavioral measures in cognitive tasks are often low dimensional (e.g., choice probability, reaction time), which makes it difficult for perturbations to resolve the precise sub-computation that may be disrupted. Here, we used rich, multifaceted measures of behavior, large-scale statistically powerful behavioral datasets and neural perturbations and recordings to richly describe cognitive strategies and identify a precise computation supported by the OFC for state inference.

## Results

### Behavioral evidence for state inference

We developed a temporal wagering task for rats, in which they were offered one of several water rewards on each trial, the volume of which (5, 10, 20, 40, 80µL) was indicated by a tone^17^ (Figure 1A). The reward was assigned randomly to one of two ports, indicated by an LED. The rat could wait for an unpredictable delay to obtain the reward, or at any time could terminate the trial by poking in the other port (“opt-out”). Reward delays were drawn from an exponential distribution, and on 15-35 percent of trials, rewards were withheld to force rats to opt-out. How long rats waited before opting out provides a robust analog behavioral measure of their subjective value of the offered water reward^14,17–19^. Rats were trained in a high-throughput behavioral training facility using computerized, semi-automated procedures to generate statistically powerful datasets across hundreds of animals^17^ (N=349 rats).

**Figure 1:**
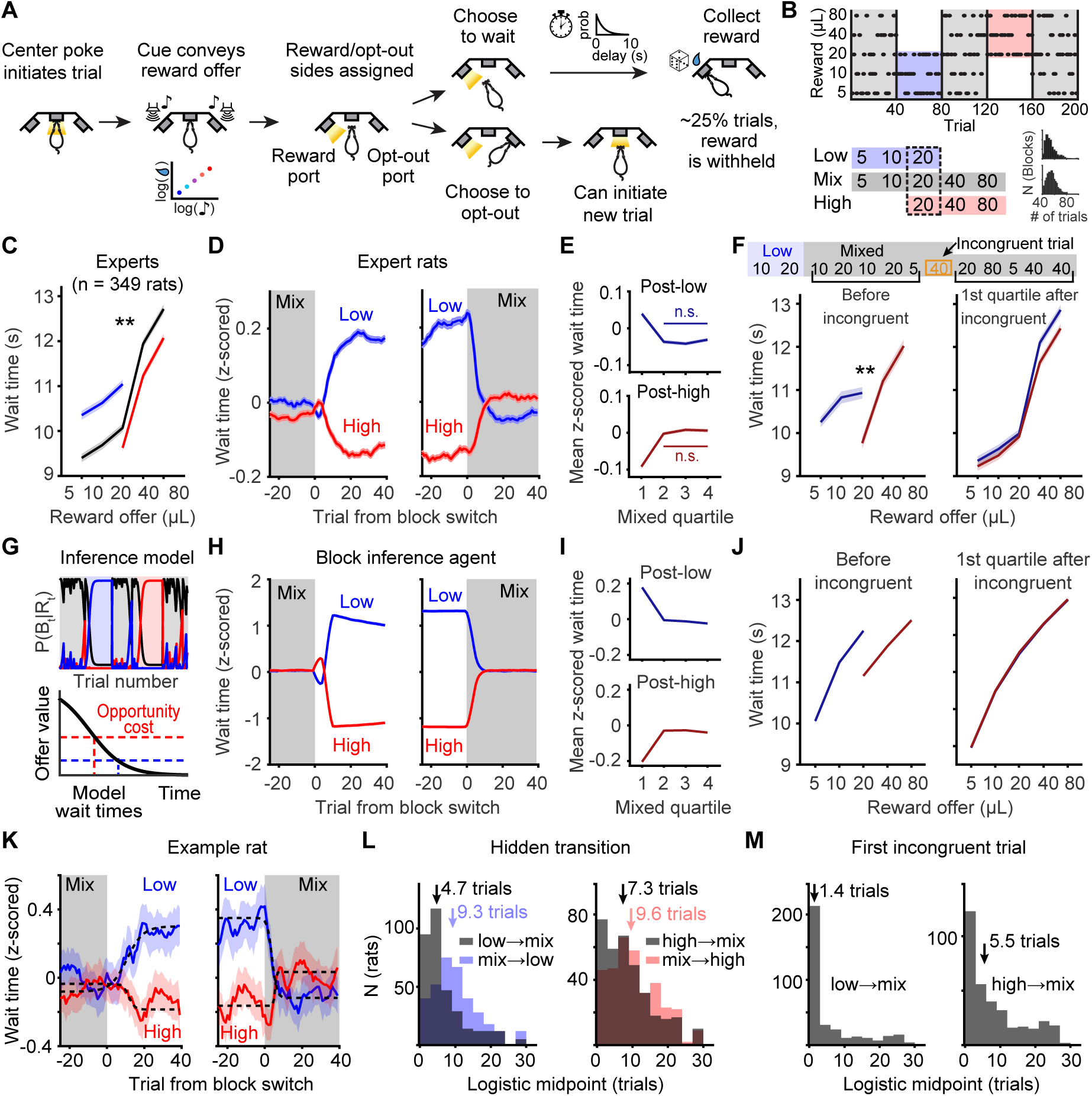
Behavioral evidence for state inference. **A.** Schematic of behavioral paradigm. **B.** Block structure of task, and histograms of experienced block durations for two example rats. **C.** Mean wait time on catch trials by reward in each block averaged across expert rats. *p* << 0.001, Wilcoxon signed-rank test comparing wait times for 20µL in high versus low blocks across rats. **D.** Mean (+/-s.e.m.) z-scored wait time at block transitions from mixed blocks into high or low blocks (left) and high or low blocks into mixed blocks (right), N = 349. Data were smoothed with a causal filter to not introduce pre-transition artifacts, by averaging with past but not future trials (5 trial window). **E.** Wait times within different quartiles of mixed blocks for expert rats. p-values for effect of quartiles 2-4 on wait times from one-way ANOVA, post-low *p* = 0.23, post-high *p* = 0.23. **F.** *left*, Wait times before the first incongruent trial in mixed blocks (*p* << 0.001) *right*, Wait times in the first quartile of mixed blocks after the first incongruent trial, which signals a block switch. Curves are conditioned on the previous block type. Bonferroni-corrected p-values for Wilcoxon signed-rank test comparing wait times conditioned on previous block type: 5µL *p* = 1, 10µL *p* = 0.76, 20µL *p* = 1, 40µL *p* = 1.67 × 10^−10^, 80µL *p* = 9.71 × 10^−10^. **G.** Model schematic of inferential agent that compares a time-varying value function to a block-specific expectation of average reward, i.e. opportunity cost. **H.** Mean z-scored wait times from a behavioral model that inferred the most likely block and used fixed, block-specific estimates of opportunity cost to decide how long to wait. **I.** Block inference model predicts that wait times should be stable within mixed blocks after a block switch has been inferred. **J.** Block inference model predicts sensitivity to the previous block type before (*left*) but not after (*right*) the first incongruent trial. **K.** Mean z-scored wait times for one example rat for each block transition, with best fit logistic functions overlaid. **L.** The distribution of x values (trials after a block transition) that corresponded to the midpoint of logistic fits for each rat, for different block transitions. Arrows show median values. **M.** Distribution of midpoint values as in panel L, but when wait times were aligned to the first incongruent trial in mixed blocks. Arrows indicate median values. Incongruent trials do not exist at transitions into high or low blocks, which is why those block transitions are not shown.

The task contained latent structure: rats experienced blocks of 40 completed trials (hidden states) in which they were presented with low (5, 10, or 20µL) or high (20, 40, or 80µL) reward volumes^17,18^. These were interleaved with mixed blocks which offered all rewards (Figure 1B). While blocks had a length of 40 completed trials, because rats prematurely broke fixation on a subset of trials, in practice block durations were highly variable (Figure 1B). The hidden states differed in their average rewards and therefore in their opportunity costs, or what the rat might miss out on by continuing to wait. According to foraging theories, the opportunity cost is the long-run average reward, or the value of the environment^20^. In accordance with these theories^20,21^, well-trained rats adjusted how long they were willing to wait for rewards in each block, and on average waited 10% less time for 20µL in high blocks, when the opportunity cost was high, compared to in low blocks (Figure 1C).

Expert rats’ wait time behavior was well-captured by an inferential strategy in which they inferred the reward block and used a fixed estimate of opportunity cost based on that state inference^17^ (Figure 1G). We next sought to characterize behavioral dynamics at block transitions. The wait times were modulated by rewards and blocks, so analyzing raw wait times would conflate both factors (Figure S1A). To isolate the effects of blocks, we first de-trended wait times to account for slow changes over the session, which were modest but present in some rats. We then z-scored the wait times for each reward independently, before averaging over z-scored wait times around block transitions (Figure 1D). As a complementary analysis, we also averaged detrended wait times for 20µL only (Figure S1B). Both analyses isolated contextual effects from the effects of reward volume on wait times, and yielded comparable results (Figure 1D, Figure S1B), although including all rewards provided greater statistical power. Notably, the inferential model captured behavioral dynamics at block transitions (Figure 1H; Figure S1C,D).

We further reasoned that an inferential strategy would produce stable within-volume wait times in mixed blocks once the animals inferred that the block had changed, as opposed to incremental adjustments based on recent rewards. To test this, for each rat, we again de-trended and z-scored the wait times for each reward, before pooling over different reward offers. We computed the mean z-scored wait times in each quartile of mixed blocks that were preceded by low or high blocks. Consistent with a state inference strategy, rats changed their behavior abruptly, within the first quartile of the mixed block, and then exhibited stable wait times over quartiles 2-4 (Figure 1E). Notably, the inferential model’s use of fixed, block-specific estimates of opportunity cost reproduced stable wait times in later portions of mixed blocks (Figure 1I).

Inferences at transitions into mixed blocks were likely driven by trials offering rewards that were not present in the previous block, which we refer to as incongruent trials (e.g., 40/80*µ*L after low blocks, 5/10*µ*L after high blocks). These trials unambiguously indicate transitions into a mixed block for an inferential agent. Rats’ wait times in the first quartiles of mixed blocks before the first incongruent trial were not different from wait times in the preceding block (post-low: p = 0.63, post-high: p = 0.35; Wilcoxon signed-rank test). How long rats waited for 20*µ*L after the hidden transition significantly differed depending on the previous block (Figure 1F *left*). These data are inconsistent with rats anticipating block transitions due to the block length. For 5/10/20*µ*L volumes, wait times *after* the first incongruent trial converged to a common value for each reward, consistent with rats rapidly inferring a transition into a mixed block following these highly informative trials (Figure 1F *right*). These patterns were predicted by the inferential behavioral model (Figure 1J). Interestingly, there was a small (∼450ms) but significant difference in expert rats’ wait times for 40 and 80*µ*L after the incongruent trial, depending on the previous block. To better understand this difference, we fit logistic functions to each rat’s wait times at transitions into mixed blocks (Figure 1K). Behavior was slower to change at transitions from high into mixed blocks, compared to low into mixed blocks, when data were aligned to the true block transition or the first incongruent trial (p = 1.56×10^−6^, p = 6.51×10^−8^, respectively; Wilcoxon signed-rank test comparing midpoints of logistic functions for individual rats; Figure 1L, M). This suggests that rats, like humans^22^, may exhibit biased or overly-optimistic belief updating: rats were quicker to infer a transition to a better block (low-to-mixed) but required more evidence to infer a transition to a worse block (high-to-mixed; Figure 1L,M), resulting in lingering high block beliefs early in the mixed block, even after the first incongruent trial (Figure 1F, *right*). This was likely apparent for large but not small reward offers, because large rewards reinforced lingering beliefs about high blocks. These data suggest that rats performed less efficient inference at transitions into relatively worse blocks, although effects were modest, with a difference of ∼4 trials (Figure 1M).

While incongruent trials indicate transitions into mixed blocks, there are no similar trials that indicate transitions from mixed into high/low blocks. Therefore, inference-based agents should show faster and sharper wait time changes into mixed blocks compared to the other blocks. We found significantly faster transitions (smaller logistic midpoints) into mixed compared to into low and high blocks (Figure 1L,M). These data build on our previous findings^17^, and suggest that rats’ behavior is well accounted for by an inferential strategy.

One potential alternative strategy that we considered was divisive normalization, in which the value of an option is divided by the sum of past rewards in some integration window^18,23^. While divisive normalization with short integration windows (∼10 trials) could produce fast behavioral changes at block transitions comparable to an inferential agent (Figure S2A-C), this model predicts trial-by-trial changes in wait times within a block (Figure S2D,E). Consistent with our previous findings^17^, we did not observe sensitivity to the previous reward (Figure S2D), and only very weak sensitivity to the sum of past 10 rewards in a given block, which could reflect mistaken inferences following sequences of large or small offers (Figure S2E). The inferential model, but not the divisive normalization model, captured these behavioral patterns (Figure S2D,E). Additionally, while the inferential model predicted faster behavioral changes into mixed blocks, compared to high and low blocks, divisive normalization with a short integration window predicted a different pattern of transition dynamics that was inconsistent with rats’ behavior (Figure S2F-I). This was true across a range of parameter values (Methods) and likely reflects the nonlinearity in the divisive normalization computation. This suggests that the inferential model better accounts for multiple aspects of rats’ behavior, including behavioral dynamics at block transitions.

### OFC inactivations impair belief updating

To determine whether OFC dynamics were causal to state inference, we performed bilateral infusions of the GABA agonist muscimol, targeted to the lateral OFC (LO; Figure 2A). Simultaneous electrophysiological recordings with Neuropixels probes confirmed that muscimol completely silenced neural activity within 1.25mm of the infusion site, indicating that our perturbations silenced LO, agranular insula and ventral OFC, but spared the medial bank of the prefrontal cortex (e.g., PL, IL, CG1; Figure S3A,B).

**Figure 2:**
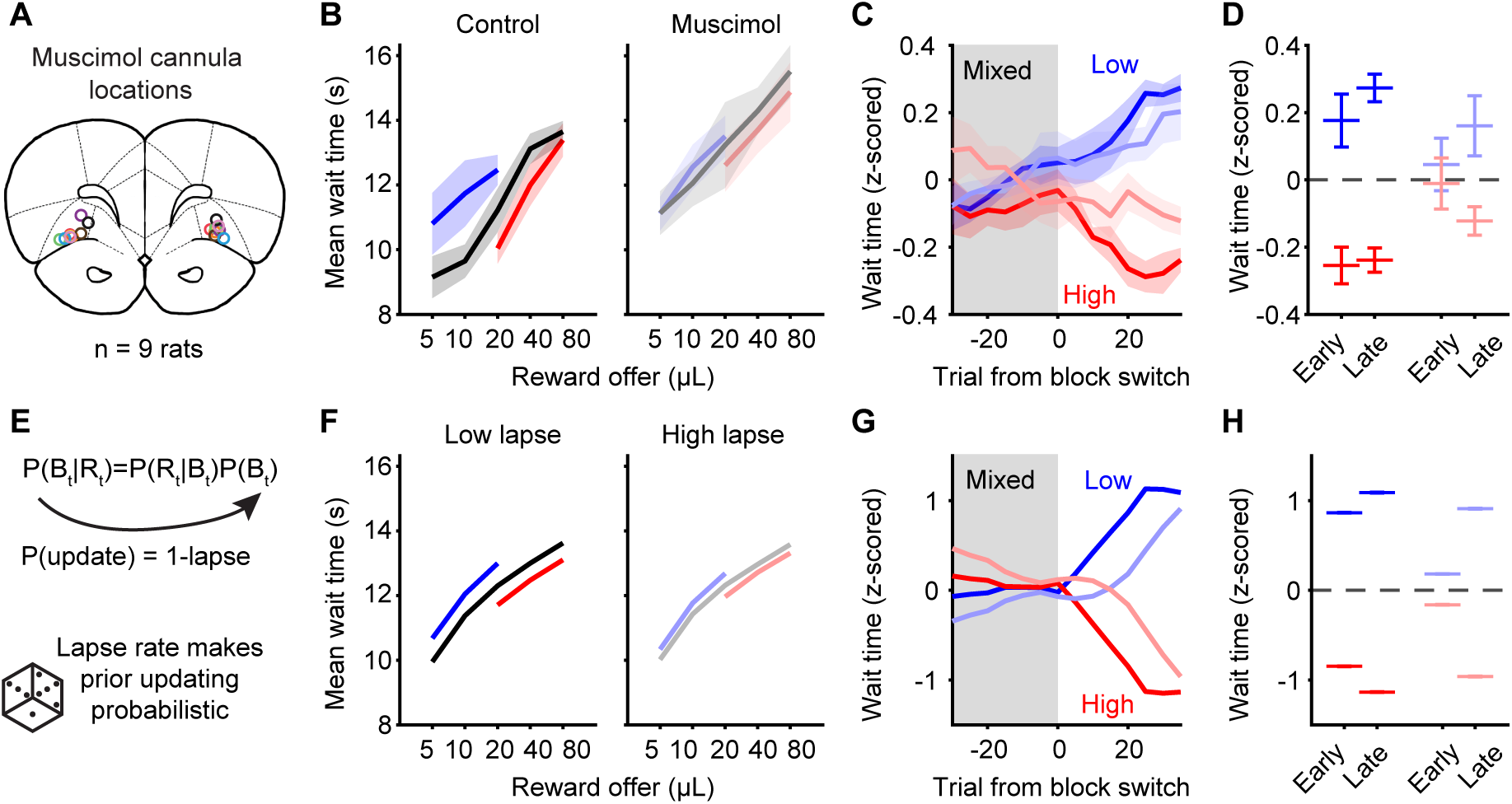
OFC supports belief updating for state inference. **A.** Location of muscimol guide cannulae in LO (N=9 rats). **B.** Mean wait times in low, mixed, and high blocks for rats in control or muscimol sessions. Muscimol produces a significant reduction in the wait time ratio, or the wait time for 20*µL* in high/low blocks (p = 0.004, Wilcoxon signed-rank test). **C.** Mean changes in z-scored wait times as animals transition from mixed into low or high blocks. Dark lines are control sessions and light lines are muscimol sessions. **D.** Mean changes in z-scored wait times early (trials 15-20) and late (trials 35-40) in a block after transitions from a mixed block. *p* = 0.004, control early; *p* = 0.004, control late; *p* = 0.65, muscimol early; *p* = 0.054, muscimol late; Wilcoxon signed-rank test. **E.** The inferential model updates its prior beliefs recursively. We introduced a lapse rate in the model which dictated a probability with which the prior was not updated, and instead remained the same for the next trial. **F.** Increasing the lapse rate (probability of the prior remaining the same) reproduced the reduction in block sensitivity observed with muscimol inactivation. **G,H.** Increasing the lapse rate made the model change its wait time behavior more slowly at block transitions.

Inactivating OFC impaired rats’ sensitivity to hidden reward states: while in control sessions, animals strongly modulated how long they waited for 20µL in each block, muscimol reduced this modulation (Figure 2B, Figure S4A-C). Moreover, OFC inactivations made rats slower to adjust their wait times following a block transition (Figure 2C). To quantify this effect, we split the high and low blocks into early and late groups of trials (see Methods). In control sessions, sensitivity to hidden reward states was significant early and late in the block, consistent with rapid behavioral adjustments based on state inference. However, in muscimol sessions, the rats were insensitive to hidden states early in the block, although by the end of the block, contextual effects were apparent (Figure 2D). These data show that inactivating OFC did not completely eliminate sensitivity to hidden states, but slowed the dynamics by which rats adjusted their behavior at state transitions.

OFC has been hypothesized to compute subjective values to drive choice^24–26^. If OFC is necessary for driving wait time behavior based on the relative subjective value of reward offers, we hypothesized that rats’ wait times should be less sensitive to the offer volume during muscimol sessions. To test this, we regressed wait times against reward volume in each block for each rat. We did not find a significant change in the slope of the wait-time curve for any block (Figure S4E), suggesting that subjective value computations were not impaired, *per se*. There was a significant increase in the offset for mixed and high blocks, but not low blocks (Figure S4D). This may reflect a systematic effect on the behavioral policy (i.e., overall willingness to wait).

OFC has been implicated in supporting goal-directed behaviors, as opposed to behaviors that are “model-free” or do not require the use of a world model^3^. Therefore, one possibility is that inactivating OFC caused expert rats to revert to an incremental, trial-by-trial strategy for estimating the opportunity cost, for instance, via divisive normalization or canonical model-free reinforcement learning^17^. An incremental strategy predicts that wait times in a given block should be sensitive to the magnitude of previous reward offers, potentially for several trials in the past. However, by several measures, rats’ wait times in mixed blocks remained insensitive to previous rewards, suggesting that they did not revert to using an incremental adaptive strategy (Figure S4F-I). Additionally, block modulation of trial initiation times, which we previously showed reflected a model-free reinforcement learning strategy^17^, was not affected by OFC inactivation (Figure S4J-L).

We next turned to the inferential model to characterize how OFC inactivations affected behavior (Figure S5A-D). The model uses Bayes’ Rule to compute the posterior probability of each block given a reward offer by combining the likelihood, or the probability of encountering the reward in a given block, with the prior belief about the block. The prior over blocks is recursively computed: the posterior on one trial becomes the prior on the next trial^17^.

We introduced a lapse rate into the model that dictated the probability with which the posterior became the prior on the next trial (Figure 2E). Increasing this lapse rate increases the probability that the prior on trial t is the prior from t-1 rather than the posterior from t-1, in other words, it makes the prior beliefs “sticky.” Increasing the lapse rate reproduced the qualitative effects of OFC inactivations, including reduced sensitivity to hidden states and slower behavioral changes at block transitions, while also producing wait times that were largely insensitive to previous rewards within a block (Figure 2F-H). In contrast, reducing the quality of the prior (Figure S5E-G), or making the block-specific opportunity costs more similar (Figure S5H-J), were unable to capture all of the effects of OFC inactivations. These results suggest that OFC supports hidden state inference by updating subjective beliefs based on outcomes.

### Single neuron correlates of state inference

We next sought to characterize neural correlates of state inference in OFC. We performed electrophysiological recordings from the lateral OFC (LO/AI) in well-trained rats using chronically-implanted Neuropixels probes (N=35 rats; Figure 3A).

**Figure 3:**
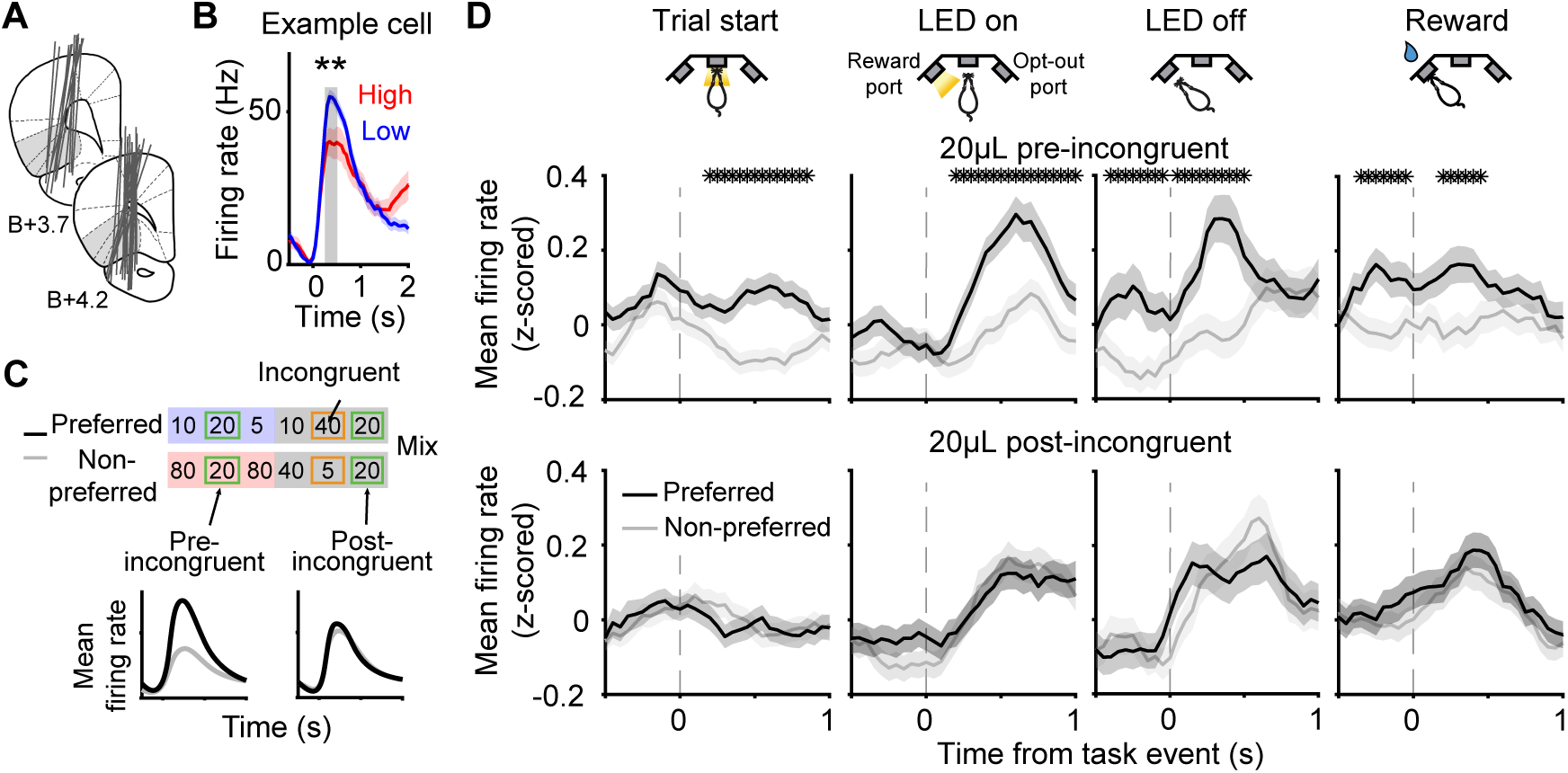
Single neurons in OFC reflect inferred states. **A.** Location of Neuropixels probes. **B.** Example neuron whose firing rate was significantly different in high versus low blocks in the window [250-500ms] after reward. **C.** Schematic of 20*µL* trials preceding versus following the first incongruent trials in mixed blocks. **D.** Mean (± s.e.m) firing rates for OFC neurons with significant block sensitivity (Trial start: 1760/5249, LED on: 1146/5249, LED off: 769/5249, Reward: 1294/5429) on 20*µL* trials preceding (upper) versus following (lower) the first incongruent trial in a mixed block, for data aligned to each task event. Asterisks indicate a significant difference in firing rates (*p <* 0.05, non-parametric permutation test).

To determine if neurons in OFC reflected inferred state transitions, we focused on trials offering 20*µ*L before and after incongruent trials following transitions into mixed blocks. As described previously, incongruent trials (which do not exist at transitions from mixed into high or low blocks) unambiguously reveal that the block has changed. If OFC dynamics reflected the inferred block, then the response to the same reward - 20*µ*L - should elicit a different firing rate before versus after the incongruent trial, because the inferred block should be different on these trials.

We first selected neurons whose firing rates were significantly different in high versus low blocks in the [0.25 0.5s] window after each task event (two-sample t-test, *p <* 0.05). We deemed the block for which they had higher (lower) firing rates the preferred (non-preferred) block for that cell (Figure 3B). Sessions without both transition types (preferred to mixed and non-preferred to mixed) were excluded. We then compared the average z-scored firing rates over these neurons for 20*µ*L trials immediately preceding versus following the first incongruent trial in a mixed block (Figure 3C). As expected, neurons exhibited different firing rates before the incongruent trial, presumably because the rat assumed it was in the neurons’ preferred or non-preferred (high or low) block (*p <* 0.05, non-parametric permutation test; Figure 3D). However, after the incongruent trial, the firing rates on 20*µ*L trials collapsed to a common value, regardless of the previous block type, consistent with rats’ inferring a transition to a common mixed block. This shows that firing rates on identical trials are different depending on whether they precede or follow trials that are informative of state transitions. This result was observed at all task events throughout the trial (Figure 3D).

Recent work in mice has argued that prior beliefs about blocks are represented in all areas of the brain, including early sensory regions^27^. To determine if recognition of incongruent trials was a ubiquitous feature of cortex, we analyzed units that were outside of LO (the Neuropixels probe also traversed through M1 and piriform cortex). We also recorded neurons from the secondary visual area V2. Neurons across all sampled areas seemed to exhibit similar coarse sensitivity to the reward blocks, as classifiers were able to decode the block identity to a comparable degree across brain regions (Figure S6A). However, neurons in M1, piriform cortex or V2 did not consistently exhibit differential firing rates on 20*µ*L trials preceding incongruent trials that collapsed to a common value following these trials (Figure S6B-D). While neurons in these areas did exhibit this pattern of activity early in the trial, at the time of the offer cue, and for M1, at the delay start, there was no neural sensitivity to inferred state transitions at the time of the reward cue after the delay, or at reward consumption, suggesting that encoding of inferred states at the time of the outcome might be unique to OFC. Overall, this indicates that persistent encoding of inferred states before and after incongruent trials was not a cortex-wide phenomenon.

We next sought to determine whether this single-cell signature of state inference was broadly distributed across the OFC population, or restricted to specific subsets of neurons. To summarize task-related responses at the single neuron level, we used a dimensionality reduction method called tensor components analysis (TCA^28^). We constructed a third order data tensor where each row corresponded to the z-scored firing rate of an individual neuron, aligned to different task events; in the z-dimension, we included the neuron’s event-aligned activity in each block. Therefore, the data tensor was organized as neurons × time × block, and the model extracted three types of factors: (1) neuron factors, which reflect how much each neuron’s activity is described by each component (i.e., loadings); (2) temporal factors, which capture time-varying event-aligned responses, and (3) block factors, which capture modulation of firing rates across blocks. TCA decomposes a third order data tensor into a sum of rank-one components. We selected the number of components based on the number at which adding additional components failed to improve the model fit^28^ (Figure S7A,B; see Methods).

We used TCA to perform unsupervised clustering of the neural responses^29^. We clustered neurons by the tensor component for which they had the maximum neuron factor or loading. Neurons that had zero loadings for all components were treated as an additional cluster (cluster 0). The temporal factors for each component captured the mean event-aligned PSTHs for neurons in each cluster (Figure 4A, Figure S7C). These data are consistent with previous findings that OFC neurons exhibit one of a relatively small subset of temporal response profiles^30,31^. We speculate that these response profiles might act as a temporal basis set for composing dynamics in the OFC. The block factors were generally flat, indicating that neurons with similar temporal response profiles likely show variable tuning for the reward blocks (Figure S7C).

**Figure 4:**
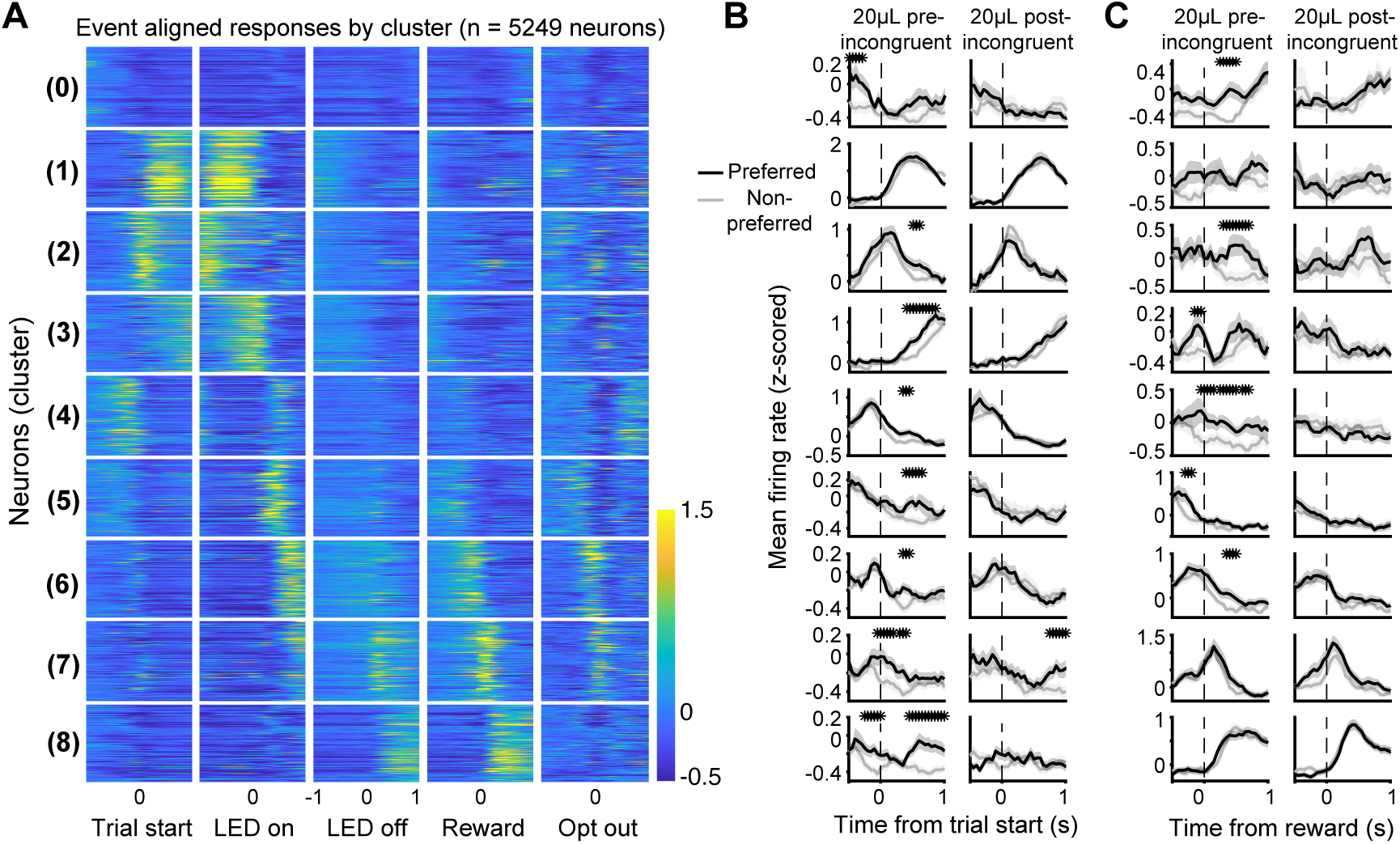
Inferred state transitions are distributed across clusters of neurons. **A.** Mean z-scored event-aligned firing rates for each neuron, sorted by the TCA component for which they have the maximum loading (see Methods). (Cluster 0, 400 neurons; cluster 1, 308 neurons; cluster 2, 494 neurons; cluster 3, 558 neurons; cluster 4, 624 neurons; cluster 5, 741 neurons; cluster 6, 857 neurons; cluster 7, 526 neurons; cluster 8, 741 neurons). **B.** Mean (+/- s.e.m) z-scored firing rates for neurons in each cluster with significant block sensitivity on 20 trials preceding (*left*) versus following (*right*) the first incongruent trial in a mixed block, aligned to trial start. Asterisks indicate a p-value *<* 0.05, non-parametric permutation test. **C.** Same as panel B but aligned to reward.

We plotted the cluster-averaged firing rates for 20*µ*L trials before and after incongruent trials. Notably, different clusters were sensitive to incongruent trials at different task events. At trial start, most clusters exhibited different firing rates before the incongruent trial that collapsed to a common value after it (Figure 4B). Other clusters encoded inferred states at the time of reward (Figure 4C), LED on, and LED off (Figure S7D,E). This suggests that sensitivity to incongruent trials at the level of the population firing rate derives from populations of neurons that are distributed across clusters, with different neurons encoding inferred states at different task events.

### Latent neural factors reflect inference

Our Neuropixels recordings generated large datasets (*>*6,000 single units). Given the scale of these data, we next sought to use dimensionality reduction to summarize task-related dynamics at the population level. Theoretical models of decision making are often described as low-dimensional dynamical systems^32,33^, so we focused on low-dimensional neural dynamics, which are also a common statistical feature of neural activity in many contexts^34–39^.

While conventional methods for extracting low-dimensional dynamics have focused on the fast (within trial) component of neural activity, a key feature of our task is that determining the value of the reward offer requires integrating over multiple timescales (e.g., evaluating the offer on single trials, inferring the reward block over many trials). To address this limitation, we developed a probabilistic hierarchical linear dynamical systems model (hLDS) that explicitly considers multiple interacting timescales. The model assumes a one-dimensional slow latent neural factor (*z_k_*) that operates at the resolution of individual trials, described by a linear gaussian stochastic dynamical system. The fast dynamics within the trial (summarized by 10 dimensional fast latent factors, *y^k^*) are assumed to operate in a similar manner. What distinguishes our approach from standard Kalman filtering is that the within trial latent dynamics are themselves dependent on the slower (evolving trial-by-trial) latent process *z_k_* (Figure 5A).

**Figure 5:**
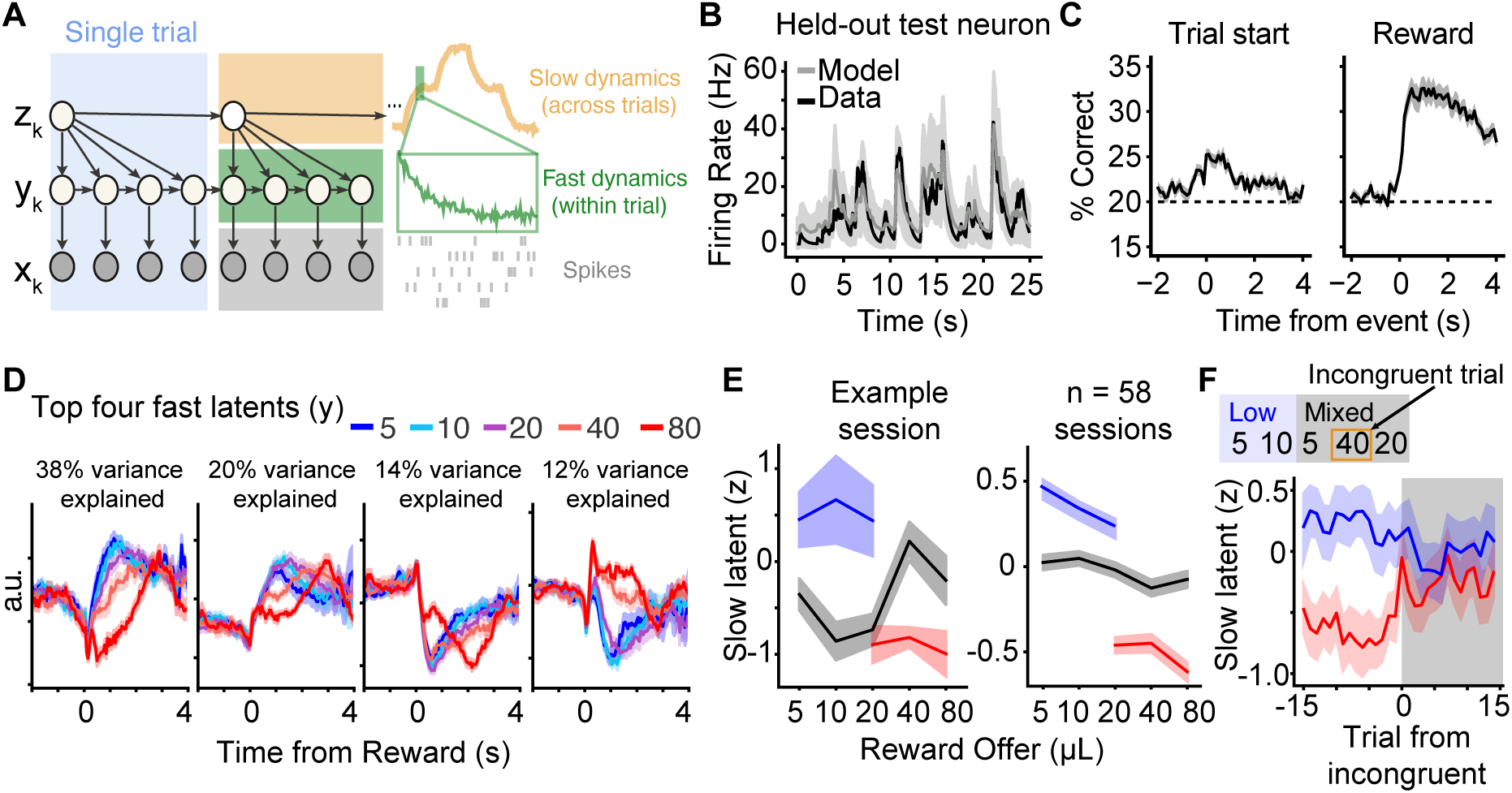
Latent factors reflect inference. **A.** Graphical model of hierarchical linear dynamical systems model (hLDS). For visualization, four fast (within-trial) latents are depicted, but the model was fit using a 1-dimensional z-latent and 10-dimensional y-latents. **B.** Model parameters fit to simultaneously recorded neurons predict the activity of a held-out test neuron. **C.** Performance of a support vector machine decoder, decoding offered reward volume from the fast latents in different time bins around the time of trial start and reward. Classifiers decoded which of five rewards were offered, so chance performance was 20% (dashed lines). **D.** The four fast latents fit to an example recording session that explain the most variance, aligned to the time of reward for trials with different reward offers. **E.** Mean slow latent on trials with different reward offers in each block for one example session (left), and n=58 recordings (right). Slow latents were z-scored for each session before combining over sessions. **F.** Schematic of incongruent trials, which unambiguously indicate a transition into a mixed block. Mean slow latent, aligned to the first incongruent trial in mixed blocks.

We fit the hLDS model to simultaneously recorded neurons using Expectation-Maximization based parameter estimation (Methods). To validate the model, we showed that it can predict the firing rates of held-out test neurons (Figure 5B), and that it better explains moment-by-moment neural responses than a dimensionality matched standard Kalman filter (Figure S8), suggesting that the hierarchical structure of the dynamics is a key feature of OFC responses during the task. Notably, model-fitting was unsupervised: the model was exclusively fit to the spikes of simultaneously recorded neurons, with no knowledge of the behavioral task. Nonetheless, the fast latents *y^k^* captured interpretable features of task-related responses, including the timing of task events and the magnitude of single trial reward offers (Figure 5C,D). It was possible to decode the reward offer from the fast latent factors, indicating that fast-timescale neural dynamics in OFC reflect reward volumes (Figure 5C). Consistent with previous results from our lab and others^19,30,31,40–42^, we found stronger encoding of reward following the outcome versus preceding choice (Figure 5C; see Discussion).

The slow latent, *z_k_*, appeared to directly reflect the hidden reward block on individual sessions (Figure 5E) and contained significant mutual information (MI) about the block (Figure 5E; MI between slow latent and blocks = 0.093, *p* << 0.001, non-parametric permutation test, Methods). We next aligned the *z_k_* latent to the first incongruent trial in each mixed block. The mean *z_k_* latent showed clear separation before the incongruent trial, and then a sharp convergence to a common value after the first incongruent trial. Therefore, rapid adjustments in latent, population-level neural factors appear to reflect changes in inferred states.

We also regressed *z_k_* against the current reward offer and the previous 10 reward offers (including an offset term) in mixed blocks. None of the previous trial coefficients were significant in these sessions, consistent with *z_k_*reflecting the inferred state within a block, as opposed to reflecting incremental reward history over trials.

### Distinct strategies over training

We next sought to characterize how behavior changed over the learning process. We analyzed the first 15 sessions during which rats were exposed to the blocks, regardless of behavioral performance. Remarkably, even in the first 15 sessions of experiencing the blocks, their wait times showed modest but significant block sensitivity (Figure 6A). However, the behavioral dynamics at block transitions appeared qualitatively different than after extensive training, suggesting a distinct psychological mechanism. Specifically, early in training, while rats showed weak changes in wait times as they transitioned from mixed into high or low blocks, behavioral changes were less apparent when they transitioned from high or low blocks into mixed blocks. Instead, the most striking behavioral feature was an offset or “DC shift” in wait times that persisted into the mixed block, possibly suggesting integration of reward history on longer timescales (Figure 6B). In contrast to expert behavior, early in training, rats’ wait times exhibited prominent within-block dynamics, suggestive of an incremental process of adjusting to the blocks (Figure 6C). Additionally, wait times in mixed blocks depended on the previous block type, both before and after the first incongruent trial, further suggesting integration of reward history on long timescales (Figure 6D). Wait times for congruent rewards in mixed blocks before the first incongruent trial were significantly different than wait times for those same rewards in the preceding block. Thus, rats modulate their wait times across latent reward blocks both early and late in training, but analysis of multiple aspects of behavior suggested distinct strategies over training.

**Figure 6:**
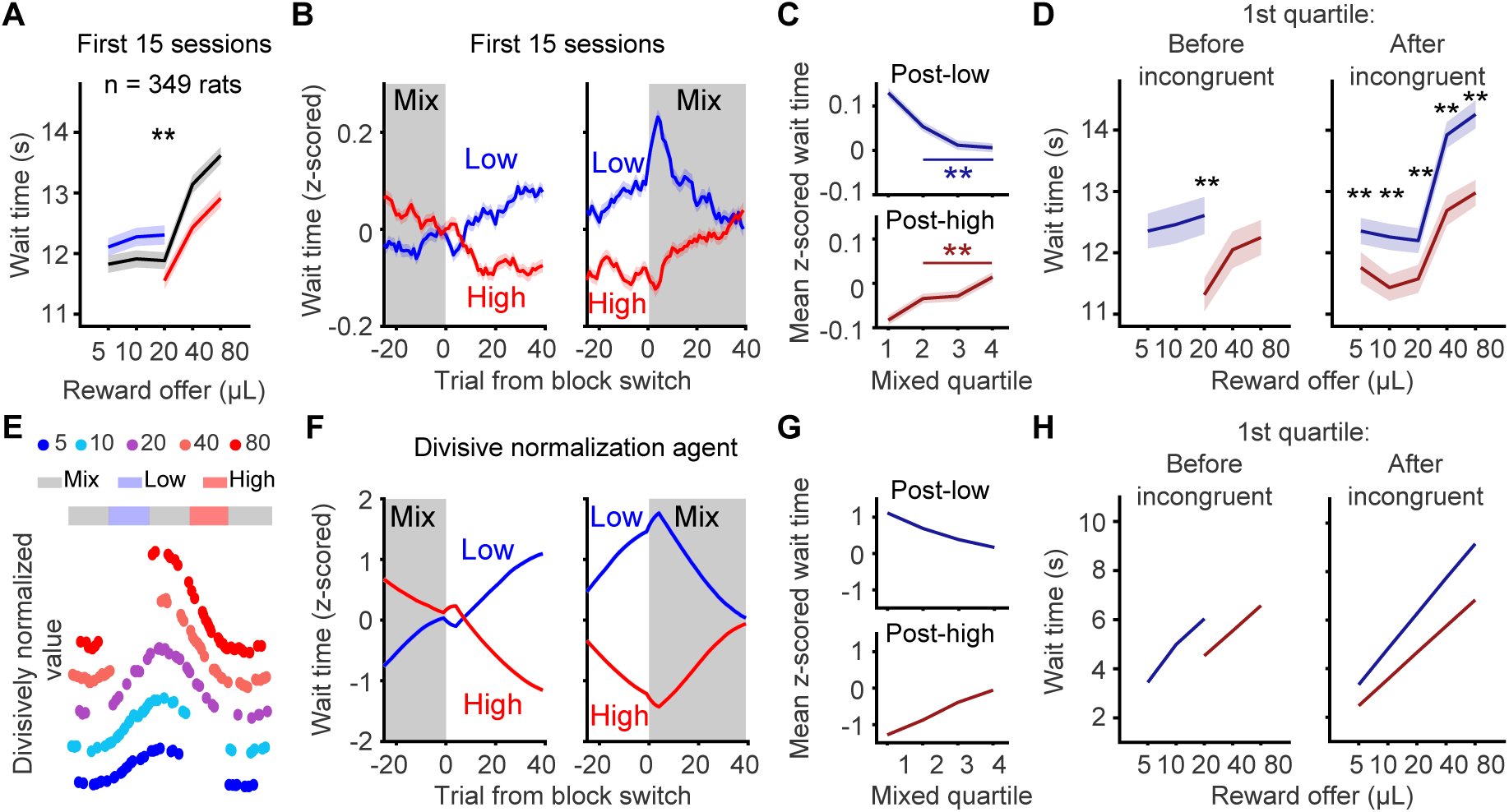
Rats use distinct strategies early in training. **A.** Mean wait time by reward in each block in the first 15 sessions of experiencing the blocks. *p* = 5.61 × 10^−19^, Wilcoxon signed-rank test comparing wait times for 20µL in high versus low blocks. **B.** Mean z-scored wait time at block transitions from mixed blocks into high or low blocks (left) and high or low blocks into mixed blocks (right), in the first 15 sessions of experiencing blocks. **C.** Mean wait times within different quartiles of mixed blocks in the first 15 sessions of experiencing the blocks; p = 0.002 (post-low), p = 0.003 (post-high); values for effect of quartiles 2-4 from one-way ANOVA. **D.** *left*, Mean wait times in mixed blocks before the first incongruent trial (*p* << 0.01). *right*, Mean wait times in the first quartile of mixed blocks after the first incongruent trial. Curves are conditioned on the previous block type. Bonferroni-corrected p-values for sign-rank test comparing wait times conditioned on previous block: 5µL *p* = 0.001, 10µL *p* = 2.1 × 10^−4^, 20µL *p* = 0.008, 40µL *p* = 7.41 × 10^−9^, 80µL *p* = 3.70 × 10^−7^. **E.** Simulated offer values of divisive normalization agent that divides the value of the current offer by the sum of previous offers in a moving window. **F.** Mean change in wait times for divisive normalization agent. **G-H.** Divisive normalization model predicts that wait times should change throughout mixed blocks, and the value of reward offers in mixed blocks depends on the previous block type both before and after an incongruent trial. All data are mean +/- s.e.m.

We considered whether early in training, rats might adopt a divisive normalization strategy^18,23^. Divisive normalization is a passive process that allows animals to adapt to different stimulus or reward distributions without requiring explicit knowledge of those distributions^43–45^. We simulated the behavior of a divisive normalization agent in our temporal wagering task with an integration window of 60 trials (Figure 6E), consistent with past work^18^ and because rats’ behavior was suggestive of long-timescale reward integration. We found that the model captured the key features of behavior early in training, including the modest behavioral changes at transitions into high and low blocks, and the prominent and sustained DC shift in wait times at transitions into mixed blocks (Figure 6F). Divisive normalization predicts incremental changes in wait times throughout the mixed block (Figure 6G), consistent with what was observed early in training (Figure 6C). Finally, within the first quartile of the mixed block, divisive normalization predicts differences in subjective values of rewards (i.e., wait times) depending on the previous block type, even after the first incongruent trial (Figure 6H). For the divisive normalization agent, the incongruent trial is no more or less informative than any other trial, so it fails to produce an abrupt change in the agent’s estimate of opportunity cost. These findings show that the divisive normalization model predicts the behavior of rats early in training, when they are naive to the blocks (i.e., “block-naive”).

Because divisive normalization is sensitive to the ordering of sequential offers, variability in the sequences of reward offers should influence the degree of block sensitivity in a session^23^. To test this hypothesis, we computed the model’s predicted wait time ratio, or the mean predicted wait time for 20*µ*L in a high block divided by a low block, and separated sessions that were in the bottom and top 50th percentiles of wait time ratios. Early in training, rats’ block sensitivity was significantly different between these groups of sessions (p=0.003, Wilcoxon signed rank test comparing wait time ratios for sessions predicted to have small or large block effects, N=349). However, in expert rats, block modulation of wait times was not different across these sessions (p=0.34, Wilcoxon signed rank test, N=349). Collectively, these data suggest that early in training, rats adapt their subjective value of rewards to the blocks via a divisive normalization algorithm (or a similar incremental, adaptive process) that integrates over long timescales (tens of trials), and that this process model can explain session-to-session variability in behavioral sensitivity to reward blocks.

### Neural dynamics reflect learning of block structure

We next recorded from the OFC of rats as they experienced the blocks for the first time. For these experiments, rats experienced extended training in mixed blocks, to isolate learning about the block structure from learning about other task parameters, such as the cue-reward mapping, catch probability, reward delay distribution, etc. When they showed consistent sensitivity of their wait times to reward volumes, they were implanted with Neuropixels probes. Rats experienced the blocks for the first time on the first recording session, and recordings were maintained for as long as units were present on the probe, sometimes for several weeks. To quantify encoding of reward offers, we computed the average discriminability index (*d*’) comparing responses on small (5/10*µ*L) versus large volume (40/80*µ*L) trials in mixed blocks. Expert rats showed more volume-encoding neurons, and higher *d*^′^ values compared to naive rats at all task events (Figure S10A,B). Consistent with decoding reward from the fast latent factors (Figure 5C), neurons reflected reward volume most strongly after the reward outcome (Figure S10B). The largest difference in *d*^′^ between expert and block naive rats was observed after reward (Figure S10B). Notably, increased reward encoding over training was specific to OFC. When we analyzed the activity of off-target neurons in motor and piriform cortex, we did not observe a comparable increase in the number of neurons significantly tuned to reward, or enhanced reward *d*^′^, in expert animals (Figure S10C-F). This further suggests that OFC’s causal contribution to state inference is reflected in unique aspects of its neural dynamics that evolve over training, and that these dynamics are not observable cortex-wide.

Pooling behavioral data across all of the recording sessions, wait times for 20*µ*L trials were not modulated by the reward block (p = 0.40, Kruskal-Wallis test for effect of block identity). However, wait times on 40 and 80*µ*L trials were significantly different between mixed and high blocks (p = 0.008, p = 0.008; Wilcoxon signed-rank test), suggesting weak contextual sensitivity (Figure 7A). As previously discussed, contextual modulation of wait times could result from inference or passive adaptation to the reward statistics. When fitting the hLDS model to simultaneously recorded neurons from block naive rats, the *z_k_* latent, on average, did not show a clear change aligned to the first incongruent trial (Figure 7B), in contrast to the experts. The mutual information between the *z_k_* latent and the blocks in the naive recordings was significantly lower than in recordings from experts (MI = 0.0491 for naive versus 0.093 for experts, Wilcoxon rank-sum p=0.0196). We again regressed *z_k_* against the current and past 10 rewards in mixed blocks. In contrast to experts, in naive recordings, the coefficient for the previous reward offer was significantly different from zero (*p* = 7 × 10^−4^), suggesting stronger representations of reward history in naive rats. Previous work has also implicated OFC in learning cognitive maps^46,47^. Given that rats were already well-trained on the task except for block transitions, and that we recorded from some rats for tens of sessions (up to 27), we hypothesized that OFC activity might reflect learning of the hidden states in some sessions.

**Figure 7:**
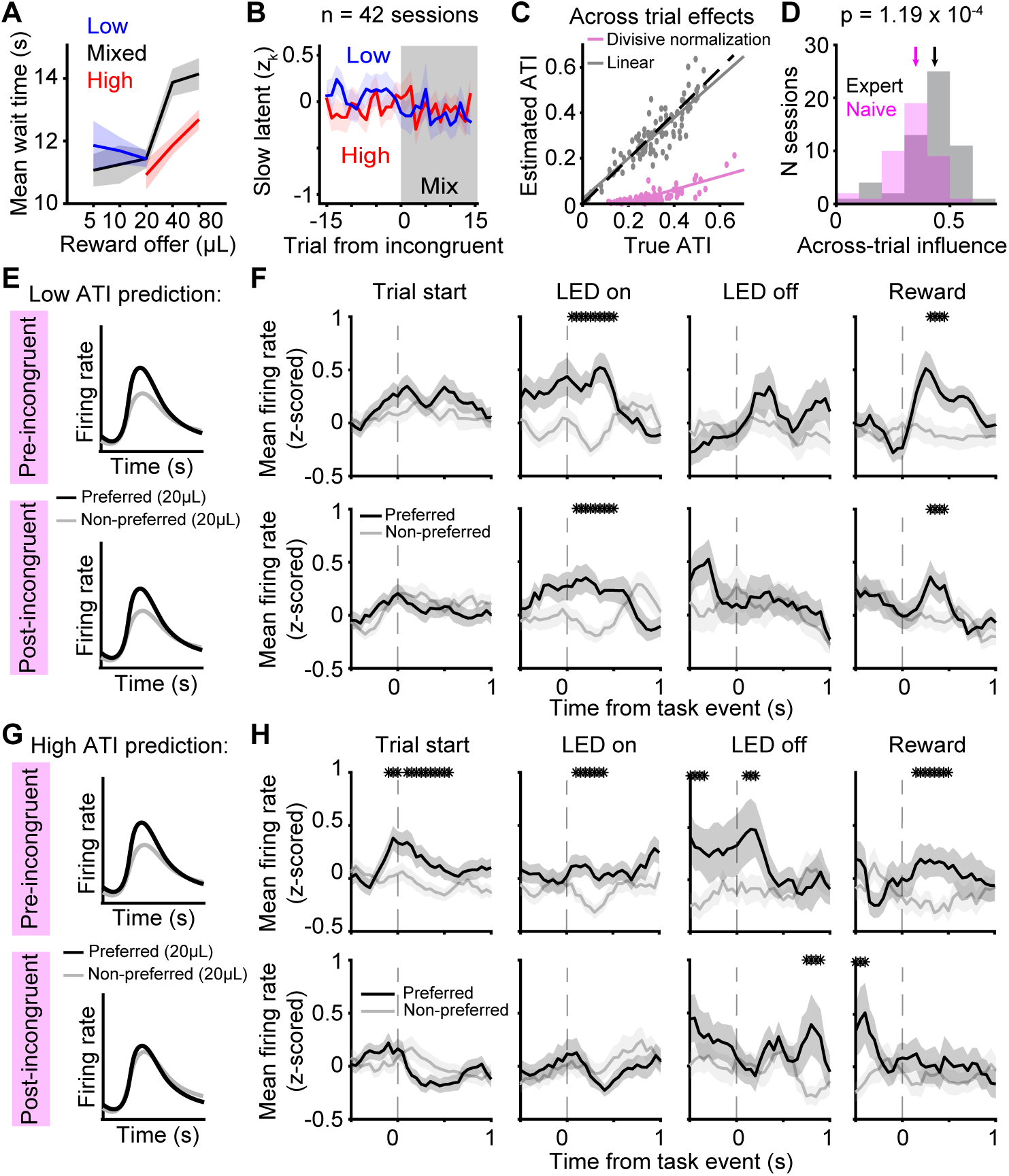
Neural activity in block naive rats reflects learning of the block structure. **A.** Mean (± s.e.m) wait time for reward volumes in each block averaged over block naive rats. Sessions were restricted to recording sessions only (96 sessions). **B.** Mean slow latent (*z_k_*), aligned to the first incongruent trial in mixed blocks. **C.** Ground truth across-trial influence (ATI) and estimated ATI for simulated population activity. hLDS estimator underestimates across-trial latents for divisive across-trial effects. **D.** Distribution of ATI metric for expert (black) and block naive (pink) sessions. Arrows indicate the median for each group (*p* = 1.19 × 10^−4^, Wilcoxon rank-sum test). **E.** Schematic of prediction for low ATI, ”naive-like” sessions. **F.** Mean firing rates for OFC neurons in low ATI sessions with significant block sensitivity on 20*µ*L trials preceding (*top*) versus following (*bottom*) the first incongruent trial in a mixed block. Asterisks indicate a p-value *<* 0.05, non-parametric permutation test. (Trial start: 274/1072 neurons; LED on: 203/1072 neurons; LED off: 93/1072 neurons; Reward: 148/1072 neurons). **G.** Schematic of prediction for high ATI, ”expert-like” sessions. **H.** Same as panel F, but for high ATI sessions. (Trial start: 224/924 neurons; LED on: 162/924 neurons; LED off: 76/924 neurons; Reward: 103/924 neurons).

It was not possible to resolve when exactly animals transitioned from divisive normalization to inference on the basis of single session behavior. However, we reasoned that the hLDS might allow us to identify sessions in which neural dynamics were more or less consistent with divisive normalization. Since the hLDS assumes across-trial computations to be *linear*, it is expected to fail at capturing across-trial dynamics that are divisive. Indeed, when simulating population activity that evolves according to inferential vs. divisive normalization across-trial updates (see Methods), we found that the hLDS estimator systematically underestimated across trial latents based on divisive normalization (Figure7C). As a concrete measure of across-trial effects on the dynamics, we computed the normalized contribution of the slow latent, relative to the total latent drive provided by both the across-trial and within trial processes - we refer to this quantity as the Across-Trial Influence or ATI. Intuitively, this metric captures how much the slow across-trial latent (*z_k_*) impacts neural firing rates on a given session. As expected, expert rats had significantly higher ATI (Figure 7D), however the distributions were overlapping, suggesting that some block naive sessions were more “expert-like”.

We hypothesized that in low ATI sessions, naive rats were using a divisive normalization strategy, but in high ATI sessions, they started to adopt inferential strategies. If that were the case, then neural correlates of inferred state transitions should differ across these sessions (Figure 7E,G). We again identified neurons whose firing rates were different in high versus low blocks, and treated the block with the higher (or lower) firing rate as preferred (or non-preferred). We compared the firing rates on trials offering the same reward, 20*µL*, immediately preceding or following incongruent trials in mixed blocks. In low ATI sessions, which we hypothesized were more divisive normalization-like, neural responses that were sensitive to the high and low blocks (at LED On and reward) remained significantly different after the incongruent trial, which is inconsistent with these rats inferring state transitions following these highly informative trials. In contrast, in high ATI sessions, responses were different before the transition at most events, and then indistinguishable at those events following incongruent trials (Figure 7H, lower row). Notably, we did not observe obvious differences in recordings from expert rats with low versus high ATI values.

## Discussion

Multiple, independent lines of evidence from behavior, perturbations, and neural recordings indicate that well-trained rats infer hidden reward states when deciding how long to wait for rewards. Neural signatures of state inference - different responses for exactly the same reward preceding versus following incongruent trials - were present at the level of single neurons, and population-level latent neural factors reflected abrupt changes following these trials. In many cognitive tasks (e.g., the two-step task), behavior on only a small subset of trials is diagnostic of different strategies^48,49^. In the limit, such as for single-shot inferences or outcome devaluation, only a single trial is used to identify or rule out particular cognitive strategies. In our task, incongruent trials were rare, typically occurring only once or twice in each recording session. We overcame statistical challenges of trial-limited analyses using a brute force approach that included high-throughput training of hundreds of rats, neural recordings of thousands of neurons in dozens of animals, and dimensionality reduction of neural data.

A major outstanding question in neuroscience, psychology and machine learning is how brains learn the states of the environment and their transition structure without supervision. In our task, the question is how the rats learn the block structure, including that there are three blocks, the reward distributions in each block, and perhaps even the average block length and alternation structure. One challenge for learning these features is that the blocks change on long timescales (tens of trials, several minutes). Another challenge is that the mixed block includes the reward sets in both low and high blocks, so changes in reinforcement between states are subtle compared to extinction, reinstatement, or reversal learning paradigms. Theories of state discovery in reinforcement learning posit a key role for large prediction errors^50^ or tonically small prediction errors^51^, but transitions from mixed into high or low blocks will not necessarily produce outlier prediction errors because of the overlapping reward distributions. While latent cause models have been developed for conditioning and extinction paradigms^52,53^, hippocampal remapping^54^, and motor learning^55^, these provide a more computational-level description and in some cases fail to specify how certain statistics in the environment are learned (e.g., context duration^53^). Empirically, we found that early in training, rats passively adapt to reward blocks by a divisive normalization strategy, whereas later in training their behavior reflects state inference. Algorithmic-level process models for structure learning are required to generate and test hypotheses about how exactly the learning process might unfold as animals experience and learn about the latent block structure, e.g., perhaps by monitoring higher order moments of the reward distributions in each block.

Divisive normalization is thought to be a canonical computation that supports efficient coding by allowing neurons to adjust the dynamic range of their firing rates to best represent stimulus or reward distributions^44^. While this algorithm is thought to reflect core features of neural circuits like inhibitory motifs and alleviate fundamental constraints of neural coding such as bounded firing rates, expert rats appear to “turn off” divisive normalization in favor of state inference. We speculate that if multiple strategies or neural systems can support behavior through different computations, then each system’s relative contribution to the expressed behavior may be determined by a winner-take-all mechanism^32^, weighted averaging^56,57^, or other arbitration process.

Previous studies in mice have found that task- or behavior-related dynamics are highly distributed and observable in most or all areas of the brain^58–61^. However, just because neural dynamics correlate with task-related variables, that does not mean that those dynamics are causal to behavior^62^. A recent study argued that prior beliefs about blocks (which dictated reward probabilities in a two-alternative forced choice task) were represented brain-wide^27^. That study employed a more permissive definition of prior beliefs that included action repetition, and they found that neural signals reflecting this term were ubiquitous, and observable even in early sensory areas. Similarly, we could decode the reward blocks to a comparable degree from neural activity in all sampled areas including V2, M1, and piriform cortex. However, we refrain from interpreting neural representations of reward blocks *per se* as reflecting computations for state inference, as these representations could reflect many factors including reward history, divisive normalization of value, or even motivation and arousal. By contrast, persistent neural encoding of inferred state transitions, controlling for encoding of reward volumes, was uniquely observed in the OFC of expert rats performing our task. This suggests that inferring hidden state transitions likely engages cognitive and neural computations that are preferentially supported by OFC. More generally, specific trials or task features that are diagnostic of particular computations may resolve more modular neural representations (i.e., dynamics that are specific to brain areas performing those computations).

While studies in primate OFC have reported strong encoding of offer values, we found more pronounced encoding of rewards following reward delivery compared to at the time of the offer cue, consistent with previous studies in rats^19,30,31,40–42^. Early in training, neural sensitivity to the blocks first emerged at the start of the delay period, when the value of the environment might influence the wait time policy, and following reward delivery, which may be the task epoch during which belief updating occurs. We speculate that perhaps because rewards are probabilistic, beliefs about latent reward states may be primarily updated following reward receipt.

A general observation about cortical responses, particularly in the frontal cortex, is that individual neurons respond to diverse combinations of task variables. Studies in the motor system have argued that single neuron heterogeneity derives from variable contributions of individual neurons to population-level latent factors that support the actual computation being performed^63^. While in motor cortex, the computations supported by neural dynamics can be reasonably assumed (e.g., motor preparation and execution), in complex cognitive tasks, there are many quantities and abstract relationships that often must be computed. Theories of mixed selectivity argue that diverse responses at the single neuron level endow downstream circuits with flexibility for decoding different variables depending on changing task demands^64,65^. However, it can be difficult to know which computations are specifically supported by the piece of tissue under study, as well as different downstream recipient circuits. Here, we demonstrated a causal relationship between recorded OFC dynamics and a precise behavioral computation, updating beliefs about hidden reward states, consistent with previous studies^1,3,13,14^. Our unsupervised analysis method revealed population-level neural factors that reflected task computations over multiple timescales, analogous to the motor system but in the context of a cognitive behavioral task. We found that these population-level factors reflected identifiable changes in tuning at the single neuron level, deriving from neurons that reflected inferred state transitions at different timepoints. Collectively, our data identify neural correlates of inference and show that these dynamics causally update belief distributions over abstract, latent states of the environment.

## Methods

### Subjects

A total of 349 male and female Long-evans rats between the ages of 6 and 24 months were used for this study (*Rattus norvegicus*). Animal use procedures were approved by the New York University Animal Welfare Committee (UAWC #2021-1120) and carried out in accordance with National Institutes of Health standards.

Rats were pair housed when possible, but were occasionally single housed. Animals were water restricted to motivate them to perform behavioral trials. From Monday to Friday, they obtained water during behavioral training sessions, which were typically 90 minutes per day, and a subsequent ad libitum period of 20 minutes. Following training on Friday until mid-day Sunday, they received ad libitum water. Rats were weighed daily.

### Behavioral training

A detailed description of behavioral training has been provided elsewhere^17^. Briefly, rats were trained in a high-throughput behavioral facility in the Constantinople lab using a computerized training protocol. They were trained in custom operant training boxes with three nose ports. Each port contained a visible LED, an infrared LED and infrared photodetector for detecting nose pokes, and the side ports contained lick tubes that delivered water via solenoid vales. There was a speaker mounted above each side port that enabled delivery of stereo sounds. The behavioral task was instantiated as a finite state machine on an Arduino-based behavioral system with a Matlab interface (Bpod State Machine r2, Sanworks), and sounds were delivered using a low-latency analog output module (Analog Output Module 4ch, Sanworks) and stereo amplifier. Each trial began with the center port being illuminated. Rats initiated the trial by poking their nose in the center point, at which time the light was turn off and an auditory cue would play.

The reward offer on each trial was cued by a tone delivered from both speakers (1, 2, 4, 8, or 16kHz). On each trial, the tone duration was randomly drawn from a uniform distribution from 800ms to 1.2s. Sound pressure was calibrated for each tone (via a gain parameter in software) so that they all matched 70dB in the rig, measured when a microphone (Bruel & Kjaer, Type 2250) was proximal to the center poke. The rat was required to maintain its nose in the center poke for the duration of sound presentation. If it terminated fixation prematurely, that was deemed a violation trial, the rat experienced a white noise sound and time out period, and the same reward offer would be presented on the subsequent trial, to disincentivize premature terminations for small volume offers. Following the fixation period, one of the side LEDs lit up indicating that port would be the reward port. The reward delay on each trial was randomly drawn from an exponential distribution with a mean of 2.5s. When reward was available, the reward port LED turned off, and rats could collect the offered reward by nose poking in that port. On 15-25% of trials, the reward was omitted. The rat could opt out of the trial at any time by poking its nose in the unlit port, after which it could immediately initiate a new trial. In rare instances, on an unrewarded trial, if the rat did not opt-out within 100s, the trial ended (“time-out trial”), and the center LED turned on to indicate a new trial.

We introduced semi-observable, hidden-states in the task by including uncued blocks of trials with different reward offers. High and low blocks, which offered the highest three or lowest three rewards, respectively, were interspersed with mixed blocks, which offered all volumes. There was a hierarchical structure to the blocks, such that high and low blocks alternated between mixed blocks (e.g., mixed-high-mixed-low, or mixed-low-mixed-high). The first block of each session was a mixed block. Blocks transitioned after 40 successfully completed trials. Because rats prematurely broke fixation on a subset of trials, in practice, block durations were variable.

To determine when rats were sufficiently trained to understand the mapping between the auditory cues and water rewards, we evaluated their wait time on catch trials as a function of offered rewards. For each training session, we first removed wait times that were greater than two standard deviations above the mean wait time on catch trials in order to remove potential lapses in attention during the delay period (this threshold was only applied to single sessions to determine whether to include them). Next, we regressed wait time against offered reward and included sessions with significantly positive slopes that immediately preceded at least one other session with a positive slope as well. Once performance surpassed this threshold, it was typically stable across months. Our analysis of expert rat behavior used this criteria to select sessions for analysis. By comparison, to examine behavior early in training, for each expert rat, we analyzed the first 15 training sessions in the final training stage when they first experience the blocks, regardless of behavioral performance.

### Training for male and female rats

We collected data from both male and female rats. Male and female rats were trained in identical behavioral rigs with the same shaping procedure (see [17] for detailed description of shaping). To obtain sufficient behavioral trials from female rats who are physically smaller than males, reward offers were slightly reduced while maintaining the logarithmic spacing: [4, 8, 16, 32, 64 *µ*L]. For behavioral analysis, reward volumes were treated as equivalent to the corresponding volume for the male rats (e.g., 16 *µ*L trials for female rats were treated the same as 20 *µ*L trials for male rats). We did not observe any significant differences between male and female rats^17^.

### Behavioral models

We developed separate behavioral models to describe rats’ behavior early and late in training. We adapted a model from [14] which described the wait time, WT, in terms of the value of the environment (i.e., the opportunity cost), the delay distribution, and the catch probability (i.e., the probability of the trial being unrewarded). Given an exponential delay distribution, we defined the predicted wait time as

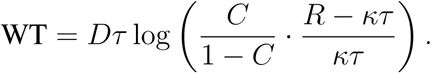

where *τ* is the time constant of the exponential delay distribution, *C* is the probability of reward (1-catch probability), *R* is the reward on that trial, *κ* is the opportunity cost, and *D* is a scaling parameter. In the context of optimal foraging theory and the marginal value theorem, which provided the theoretical foundation for this model, each trial is a depleting “patch” whose value decreases as the rat waits^20^. Within a patch, the decision to leave depends on the overall value of the environment, *κ*, which is stable within trials but can vary across trials and hidden reward states, i.e., blocks.

The inferential model has three discrete value parameters (*κ*_low_*, κ*_mixed_*, κ*_high_), each associated with a block. For each trial, the model chooses the *κ* associated with the most probable block given the rat’s reward history. Specifically, for each trial, Bayes’ Theorem specifies the following:

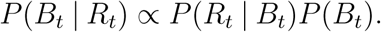

where *B_t_* is the block on trial *t* and *R_t_* is the reward on trial *t*. The likelihood, *P* (*R_t_* | *B_t_*), is the probability of the reward for each block, for example,

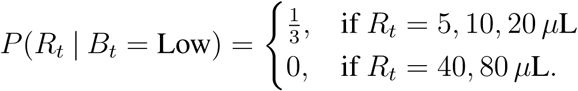

To calculate the prior over blocks, *P* (*B_t_*), we marginalize over the previous block and use the previous estimate of the posterior:

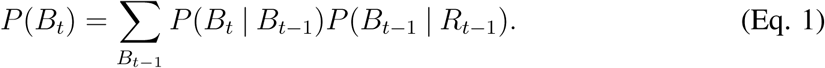

*P* (*B_t_* | *B_t_*_−1_), referred to as the “hazard rate,” incorporates knowledge of the task structure, including the block length and block transition probabilities. For example,

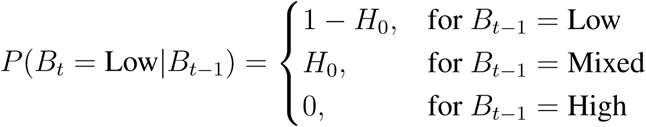

where *H*_0_ = 1*/*40, to reflect the block length. Including *H*_0_ as an additional free parameter did not improve the performance of the wait time model evaluated on held-out test data in a subset of rats (data not shown), so *H*_0_ was treated as a constant term.

### Inferential model with lapse parameter

The optimal inferential model used a recursive updating procedure in which the posterior on trial *t*_1_ became the prior on trial *t*. For the model with a lapse rate (*lapse*), this updating procedure was probabilistic such that,

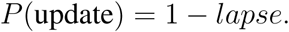

For the low lapse condition, *lapse* was set to 0. For the high lapse condition, *lapse* was set to 0.75.

### Inferential model with lambda parameter

To account for potentially sub-optimal inference across rats, we developed an inferential model with a parameter, *λ*, that generates the sub-optimal prior by weighting between the true, optimal prior (P(Bt), Eq. 1), and an uninformative prior (*Prior_nonflat_*; mixed 1/2, low and high 1/4), that is,

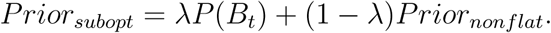

When *λ* = 1, this model reduces to the optimal inferential model, and when *λ* = 0, this model uses a non-flat prior and the block probabilities are driven by the likelihood. The *λ* parameter was set to 1 for the optimal prior case and 0.25 for the sub-optimal prior case.

### Divisive normalization model

The divisive normalization model divides the value of each offer by the sum of past rewards in some window of trials, following [18]. We modeled the wait times as being directly proportional to this term:

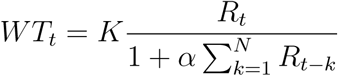

where *R_t_*is the reward offer on trial *t*, and *N* dictates the number of previous rewards, and *K* and *α* are model parameters. Previous behavioral studies^18^ suggested that dynamic valuation in humans was well-captured with an *N* of 60 previous trials, and this parameter reproduced multiple features of rat behavior early in training. For model simulations, we set *K* = 50 and *α* = 0.15.

When simulating the inferential and divisive normalization models, we treated *R_t_* as *log*_2_(*R_t_*), to be consistent with our previous studies^17^, which assumed that rats exhibited compressive utility functions^66^. However, all of our results qualitatively held if we did not log transform the reward offers (data not shown). We simulated the model using trials for which we also had rat data (non-violation trials). At the start of each new session, the trial integration window was reset, and the sum was set to the empirically determined average per-trial offer.

To quantify the transition dynamics of the divisive normalization model, we simulated the model on the sequences of trials that a subset of rats (N=50) experienced. We then computed the mean z-scored wait times in different trials around block transitions, and fit logistic functions to those curves. Fixing the integration window to 10, we varied *α* across a range [.15 .3 .45] and found that these parameters all made identical predictions about the transition dynamics, i.e., *α* does not contribute to transition dynamics. Varying the integration window did impact the transition dynamics, but we only included simulations with an integration window of 10 trials because this was the parameter that produced fast behavioral changes most consistent with expert rats’ behavior.

### Statistical analyses

Exact p-values were reported if greater than 10^−20^. For p-values smaller than 10^−20^, we reported *p* << 0.001.

### Detrending wait times

Some rats exhibited modest, gradual increases in wait times over the course of the behavioral session. To account for this, and remove potential effects of motivation and satiety from the wait times, we performed the following detrending procedure for all wait time analyses. First, we removed outlier wait times that were one standard deviation above the pooled-session mean. We fit a regression line to the mean wait time as a function of trial number, averaging over all sessions, using Matlab’s regress.m function. We then subtracted the regression line from the wait times on each session, adding back the offset parameter so that wait time units were in seconds and not zero-meaned.

### Wait time sensitivity to reward blocks

For all analyses, we removed wait times that were one standard deviation above the pooled-session mean. Without thresholding, the contextual effects are qualitatively similar, but the wait time curves are shifted upwards because of outliers that likely reflect inattention or task disengagement^17^. When assessing whether a rat’s wait time differed by blocks, we compared each rat’s wait time on catch trials offering 20 *µ*L in high and low blocks using a non-parametric Wilcoxon rank-sum test, given that the wait times are roughly log-normally distributed. We defined each rat’s wait time ratio as the average wait time on 20*µ*L catch trials in high blocks/low blocks.

### Block transition dynamics

To examine behavioral dynamics around block transitions, for each rat, we detrended the mean wait time over the course of the session. These effects were modest but in some rats, produced a slight increase in wait times over the session. We z-scored wait-times for all catch trials. We then z-scored wait times for each volume, relative to the average z-scored wait time for that volume, in each time bin (trial relative to block transition), before averaging over all volumes. We smoothed the average curve for each rat using a 5-point causal filter, before averaging over rats.

To quantify differences in wait time dynamics at different block transitions, we fit a 4-parameter logistic of the following form:

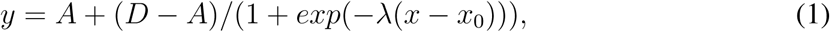

where A and D correspond to the asymptotes, *λ* is the inverse temperature parameter, and *x*_0_ is the x-value of the sigmoid’s midpoint. Statistics were performed on the best fit *x*_0_ parameter across rats. We used MATLAB’s constrained minimization function, fmincon, to minimize the mean squared error between the sigmoid and the wait time dynamics. 50 random seeds were used in the parameter search for each rat; parameter values with the minimum squared error were deemed the best fit parameters.

### Mixed block quartile analysis

To compute the mean wait times in each quartile of mixed blocks, we first detrended the mean wait time over the course of the session. We regressed mean wait time against trial number pooling over sessions, and subtracted the model-predicted effect of trial number from the wait times of each session. We then z-scored wait-times for opt-out trials of each volume separately in order to control for reward volume effects. We then separated mixed blocks depending on whether they were preceded by a low or high block. We divided each block (including violation trials) into four equally spaced bins of trials. Blocks that were fewer than 40 trials (e.g., if the rat did not complete the block at the end of the training session) were excluded from analysis. We then averaged the z-scored wait times in each quartile/bin for mixed blocks that were preceded by low and high blocks. To determine if there was an effect of mixed quartile on the wait times (i.e., if there were within-block dynamics of wait times), we performed a one-way ANOVA. Because we expected the wait times to change following an inferred state transition in the first quartile, we restricted this analysis to the second through fourth quartiles.

To characterize the mixed block wait times in the first quartile after the first incongruent trial, we separated mixed blocks depending on whether they were preceded by a low or high block, and divided each block (including violation trials) into four equally spaced bins of trials. We analyzed trials in the first bin/quartile only, and excluded trials preceding and including the first incongruent trial. We then plotted the mean wait times as a function of reward offers for trials in the first mixed block quartile after the first incongruent trial, separately for blocks preceded by low or high blocks. We compared wait times for each reward following a low versus high block using a Wilcoxon signed rank test. To correct for multiple comparisons, we multiplied each p-value by the number of comparisons (five, one for each reward). The p-values reported in the figure legend reflect this Bonferroni correction.

### Trial history effects

To assess wait time sensitivity to previous offers (Supplementary fig. 2D-F), we focused on 20 *µ*L catch trials in mixed blocks only. To minimize the impact of mistaken inferences in mixed blocks^17^, we excluded trials in which the inferential model mistakenly inferred that it was in a high or low block. We z-scored the wait times of 20 *µ*L mixed block catch trials separately. Next, we averaged wait times depending on whether the previous offer was greater than or less than 20 *µ*L. We defined the sensitivity to previous offers as the difference between average wait time for trials with a previous offer less than 20 *µ*L and trials with a previous offer greater than 20 *µ*L. We compared wait time sensitivity to previous offers across rats using a paired Wilcoxon signed-rank test.

We also assessed wait time sensitivity to the sum of several previous offers (Supplementary fig. 2G-I). We first computed the sum over the previous 10 rewarded trials for each 20 *µ*L catch trial in mixed blocks. To minimize the impact of mistaken inferences in mixed blocks, we excluded trials in which the inferential model mistakenly inferred that it was in a high or low block (although we note that rats’ actual mistaken inferences are unknown). We z-scored the wait times on these trials separately. We next averaged the z-scored wait times depending on whether the sum over the previous 10 trials was in the bottom or top 50% of sums for each rat. We compared wait time sensitivity to previous offers across rats using a paired Wilcoxon signed-rank test.

### Trial initiation times

We excluded trial initiation times above the 99th percentile of the rat’s cumulative trial initiation time distribution pooled over sessions. Ratios were computed as the ratio between trial initiation times, pooled over all volumes, in high blocks/low blocks.

### Surgical procedures

Rats were anesthetized using 3-4% isofluorane in oxygen at a flow rate of 2.5 L/minute for induction and 1-2.5% isofluorane in oxygen as maintenance for the duration of the procedure. Rats were placed in a steoreotaxic frame (Kopf instruments). 1-2 bone screws (Fine Science Tools) were inserted anterior or posterior to the target site. For muscimol experiments, guide cannulae (PlasticsOne) were then inserted bilaterally targeting AP +4.0, ML ±2.5 to a depth of 2.9mm ventral from the brain surface. Craniotomies were sealed with silicone elastomer (Kwiksil, WPI). Metabond (Parkell) was applied to the brain surface and cannulae were fixed in place with a combination of Dentin (Parkell) and dental acrylic. A custom 3d printed headcap was sealed to the skull using the same mixture to protect the implant. Guide cannulae were covered with a dustcap and dummy insert extending 0.5mm below the surface of the guide. Animals restarted training after 5-7 days to allow for recovery.

For electrophysiolgy recordings, Neuropixels 1.0 probes (IMEC) were mounted on custom 3-D printed probe mounts following [67] prior to surgery. The anesthetization procedure was the same as for muscimol rats. Before surgery, probes were dipped in the lipophylic dye DiI.

A bone screw was inserted over the olfactory bulb. Small craniotomies (∼1mm or less were drilled over OFC or cerebellum (or posterior parietal cortex). Metabond was used to coat the skull. A silver wire (A-M systems) was inserted into the craniotomy over cerebellum to a depth of approximately 1mm into the brain for grounding. The ground craniotomy was sealed with Kwiksil, and dentin was used to secure the wire to the skull. The mounted Neuropixels probe was then lowered into the craniotomy over OFC, targeting AP +3.7 and ML ±2.5, counterbalancing between right and left. Probes in secondary visual cortex (V2) were implanted at AP -4.7, ML ±4.0. Probes were lowered so the base of the probe mount sat on the skull (5.5 - 7 mm DV). The probe craniotomy was sealed with Dowsil (Dow). Puralube (Dechra) was then used to seal the space around the probe mount to prevent any liquid from getting into the chamber. The implant was then secured to the skull with Dentin and dental acrylic. Bubble wrap was placed around the implant to protect from damage. Animals were allowed to recover for at least 5 days before recording.

### Muscimol infusions

During infusions, rats were anesthetized with 1.5-2% isofluorane in oxygen at a flow rate of 2.5 L/minute. Dummy cannula were removed and cleaned with 70% ethanol. A Hamilton syringe connected to the infuser by silicone tubing filled with mineral oil was used to infuse 300-320 nL of muscimol (0.125 *µ*g/*µ*L) bilaterally over a 90s period. The infusion cannula extended 1.5mm below the end of the guide cannula. Visual confirmation of a drop in the meniscus was used to ensure successful infusion. Animals were run after a 30-45 minute recovery period. On control sessions, animals were similarly anesthetized but did not receive an infusion of muscimol. Control and muscimol sessions were alternated. Animals were given a two day “wash-out” period to prevent lingering effects of either isofluorane or muscimol. Data for those sessions was not included.

To verify inactivation of neural activity, in an acute experiment, an animal was anesthestized with isoflurane, and a Neuropixels 1.0 probe was lowered at an angle in the same craniotomy as the infusion cannula. Recordings were performed before, during, and up to 30-40 minutes after infusion of 300 nL of muscimol. Based on reconstruction of the probe track from post-mortem histology, we estimated the locations of different recording channels relative to the infusion cannula. We found robust inactivation of neural activity, relative to pre-infusion baselines, up to 1.25mm from the infusion site.

### Histology

Histology was performed on all implanted rats after experiments concluded. The tips of all infusion cannulae were within the borders of LO for all rats except 1, which was excluded from analysis. Neuropixels probe tracks were reconstructed from post-mortem histology, and the location of individual recording channels relative to areal boundaries was estimated. Channels that were estimated to be outside of LO or agranular insula (AI) were excluded from further analysis, unless otherwise stated. Cells recorded from channels estimated to be ventral to LO or AI were considered piriform cortex cells. Cells recorded from channels estimated to be dorsal to LO or AI were considered motor cortex cells. For V2 recordings, cells estimated to be outside V2 were excluded.

### Processing of neural recordings

Data were acquired using OpenEphys. Spikes were sorted by Kilosort2.0, and manually curated in Phy. Units were further curated using a custom Matlab script. Units with greater than 1% inter-spike intervals less than 1 ms, firing rates less than 1 Hz, or were completely silent for more than 5% of the total recording were excluded. To convert spikes to firing rates, spike counts were binned in 50 ms bins and smoothed using Matlab’s smooth.m function.

### Single neuron analysis of incongruent trials

Only sessions with at least 5 simultaneously recorded neurons and at least 1 full block of each type (low, mixed, high) were included. To identify neural correlates of inferred state transitions, we first selected neurons that exhibited significantly different firing rates between high and low blocks, in the [0.25 0.5s] window aligned to the time of reward delivery (two-sample t-test, p*<*0.05). For these neurons, the block that produced the higher firing rate in that window was deemed the preferred block, and the block that produced the lower firing rate was non-preferred. Pre-incongruent 20*µ*L trials were defined as the last non-violation 20*µ*L trial before an incongruent trial, regardless of whether this occurred in a mixed block or the previous low or high block. These were required to be within 10 trials before the incongruent trial, including violation trials. Post-incongruent 20*µ*L trials were defined as the first non-violation 20*µ*L trial in a mixed block after an incongruent trial. These were required to be within 10 trials after the incongruent trial, including violation trials. The first incongruent trial was defined as the first trial offering a volume not present in the previous block (5 or 10 *µ*L after a high block, 40 or 80 *µ*L after a low block) and could be any trial type (rewarded, opt out, violation). To determine if neurons exhibited different firing rates on these trials we performed a non-parametric permutation test on each 50ms time bin. We generated null distributions on differences in firing rates over groups of trial types by shuffling the labels of neurons as belonging to pre-incongruent trials or post-incongruent trials, recomputing differences in firing rates from randomly drawn groups, and repeating that procedure 1000 times. We then used this null distribution to calculate a p-value for the observed differences in firing rates between groups of rats: the area under this distribution evaluated at the actual difference of firing rates (between trial types) was treated as the p-value. To avoid false positives, we required at least three consecutive time bins to have p *<* 0.05, in order to be considered significant.

### Tensor components analysis

To fit the TCA model, we used software from^28^ https://github.com/ahwillia/tensortools and^68^ https://github.com/kimjingu/nonnegfac-matlab. Tensors were constructed by first z-scoring each neuron’s firing rate. Block responses were averaged over all volumes within a window of [-1 1s] around each trial event. Each neuron’s responses were concatenated across the 5 trial events (trial start, side LED on, side LED off, reward, optout out). Responses were then concatenated over neurons to create a 3d neurons x time x blocks tensor. Only neurons from sessions with all three block types were included. Models were fit using non-negative tensor factorization (Canonical Decomposition/PARAFAC). To initially determine the dimensionality, or rank, that should be applied to each model, we iteratively tried different numbers of dimensions, or ‘tensor components’, and computed the reconstruction error between the model prediction and data. We identified the inflection point, or the point at which adding additional components failed to reduce reconstruction error. Using all of the recorded neurons in each group of animals (expert and naive rats), these error plots suggested that the data were well-captured by a rank 8 model. Adding more than 8 components tended to yield components with flat temporal factors and negligible or zero neuron factors, suggesting that the model was overparameterized.

We grouped neurons based on the component for which they had the highest neuron factor or loading. A subset of neurons in each group had zero loadings for all components. This was because their z-scored firing rates were suppressed throughout the trial, and non-negative TCA failed to capture their task-modulation. We included these neurons as “Cluster 0” in both groups of rats (Fig. 3d-g).

### Block decoding

We used MATLAB’s multinomial logistic regression function (mnrfit) to decode the reward block from neuron firing rates [0 0.5s] after reward. We performed 5-fold cross-validation to evaluate decoder performance. Sets of trials used in each training set were balanced across both blocks and volumes. Sessions without all 3 reward blocks or fewer than 5 cells were excluded.

### Discriminability index

To measure neuronal discriminability for reward we computed the mean difference in the smoothed firing rate on small volume mixed block (5/10*µ*) trials and large volume mixed block (40/80 *µ*L) trials divided by the square root of their mean variance:

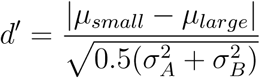

Because we computed the absolute value of the difference in firing rates, we subtracted the mean shuffled d’, computed from shuffling the data 15 times. d’ was computed in 50 ms bins, as this was the bin-width used for computing the firing rates.

### Hierarchical latent dynamical system model

The hierarchical linear dynamical systems (hLDS) model assumes a one-dimensional latent factor **z***_k_*that operates at the resolution of individual trials, with linear gaussian dynamics:

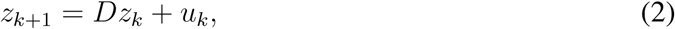

where *D* is a parameter determining the time scale of the slow dynamics and *u_k_* is independent gaussian white noise, *u_k_*∼ N (0*, σ*^2^).

The latent dynamics within a trial, **y***^k^* (of dimensionality *d*) also have linear gaussian dynamics, driven by the slow component *z^k^*:

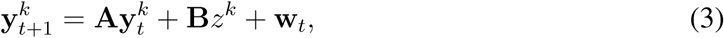

where *k* and *t* index the trial, and the time bin within the trial, respectively; the noise **w***_t_* is drawn i.i.d. from a zero mean multivariate normal distribution with isotropic variance, 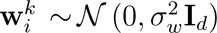. The fast dynamics are parametrized by matrix **A** and noise variance 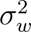 ; vector **B** parametrizes the influence of the slow latent onto each dimension of the fast dynamics. Note that the noise processes are essential for avoiding degenerate dynamics; in the absence of noise, the latents will either shrink to zero or explode with an exponential rate^69^.

Given the fast dynamics, the neural observations are conditionally independent linear gaussian:

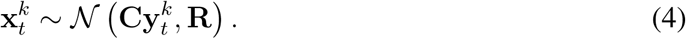

Parameter matrix **C**, of size *n* × *d* (where *n* is the number of simultaneously recorded neurons) determines the degree to which individual neural responses are affected by the low-dimensional population dynamics, with observation noise parametrized by **R**. Consistent to previous literature, we use square-root transformed binned spike counts as the neural measurements, to closer match the marginal gaussianity assumptions of the model^35,70^.

We derive filtering and smoothing updates as hierarchical analogues of Kalman filtering and smoothing. Parameters of the model were estimates maximum likelihood, via expectation maximization^71^. While we use smoothing for the E step of parameter estimation, when investigating the coding properties of the resulting latents, inference was done via filtering so as to ensure the causal temporal structure is maintained.

The model was fit independently to each session, with a fixed fast latent dimensionality of 10. We reparameterized the latent space by a well-established orthonormalization procedure^35^ that uses singular value-decomposition of the learned observation matrix C to produce an equivalent parameter set with the same observations but factorized fast latents; we sort the resulting latents based on how much data variance they explain. As the sign and magnitude of the slow latent on any given session was arbitrary, these were z-scored across sessions and signed so the mean low-block slow-latent was positive.

To evaluate model predictions for held-out test neurons, we used a standard procedure for evaluating latent state space models^38^. The model assumes that the fast latents drive neural activity via the n x d parameter C (where n is the number of neurons and d is the number of fast latents). The held-out neuron’s data was included during the fitting procedure, such that an n x d matrix C was learned. For the hold-out test, the row of C corresponding to the held-out neuron was omitted, yielding an estimate of the d-dimensional latent space y using only the n-1 neurons. That is to say, the inference procedure was identical, except using an (n-1) x d = C’ and data for all neurons except the held-out neuron. The activity for the left out neuron was then estimated by projecting this inferred latent space y back into the observation space x using the weights from the row that was left out during inference. This procedure was executed on held-out test data that was not used for fitting.

### Across Trial Influence

We defined the Across Trial Influence (ATI) metric in terms of the relative norm of the input drive from *z^k^* compared to the norm of the total change in the within-trial latents *z^k^*

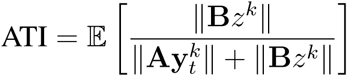

where the expectation is taken over time bins and trials. In simple terms, this quantifies how strong are the effects of the across trial latents in terms of explaining neural population responses. The ATI approaches zero when the slow latents have negligible effects on the dynamics and it approaches 1 for a scenario where there are no additional within-trial dynamics (**A** = **0**) so that the fast latents (and the neural responses themselves) are temporally i.i.d. gaussian fluctuations around a *z^k^*-determined mean.

To demonstrate the utility of the ATI for disambiguating between belief updating (i.e. additive slow latents) and divisive normalization (nonlinear coupling of across and within trial latents), we simulated data with known ground truth parameters from both types of dynamics (100 datasets each, of the same size as the actual data), and investigated the estimated hLDS parameters and associated ATI estimates. For the “expert-like” scenario, the true data generating process followed the hLDS generative model, whereas for the “divisive normalization” scenario we introduced an additional nonlinear step, directly reflecting the computations of the corresponding behavioral model:

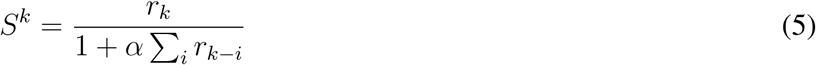

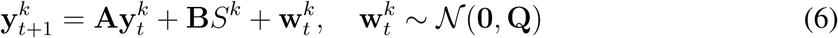

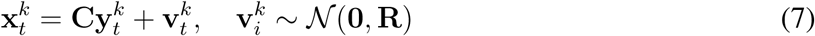

where parameter *α* and the reward integration horizon are the same as for behavior.

The ground truth ATI for the expert-like data is defined as before. In the case of divisive normalization, the ground-truth ATI is defined as:

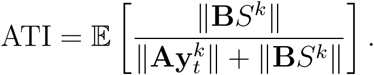

which maintains a consistent interpretation of the ratio while taking into account the different nature of the *z*’s effect on **y***_t_*. Given this simulated data, we then used an identical hLDS estimation procedure as used for the real data and compared the ground truth vs. the estimated ATIs.

### SVM decoder

We constructed a support-vector machine (SVM, using the scikit-learn library in Python) to decode reward volumes from the fast latents extracted from the hLDS, **y***^k^*. The decoder was trained and tested using trials from all blocks. Fast-latents were discretized into 250 ms time bins. We trained and cross-validated the SVM using a 10-fold cross-validation. The decoder was trained to decode 5, 10, 20, 40, or 80 muL, and trials were balanced across groups, so chance performance was 20%.

### Mutual information

To determine the relationship between the slow latent process with latent reward blocks across groups of animals, the slow-latent values were estimated for 58 sessions from expert animals and 42 sessions from naive animals. Mutual information values were computed using a non-binning MI estimator for the case of one discrete data set (block, 3 possible values) and one continuous data set (z-latent)^72^. We computed significance by shuffling the block labels across n=1000 repetitions, generating a null distribution for our test statistic with a significance threshold of *α* = 0.05. textcolorblue Mutual information between the slow latent process and blocks in expert animals (mean MI: 0.0931 bits, std: 0.0861 bits) was greater than the mutual information between the slow latent process and blocks in naive animals (mean MI: 0.0491 bits, std: 0.0557 bits) across all sessions (Wilcoxon rank-sum p=0.0196).

### Regressing slow latent against reward history

To determine if the slow latent reflected reward history, we first z-scored this variable so that magnitudes were comparable across sessions. We then regressed it against the current reward offer and the previous 10 reward offers (including an offset term), using the built in OLS regression method in the Python statsmodels package. We evaluated the significance of each regression coefficient by the t-statistic. While none of the previous trial coefficients were significant in recordings from expert animals, in naive recordings, the coefficient for the previous reward offer was significantly different from zero (*p* = 7 × 10^−4^), suggesting stronger representations of reward history in naive rats.

## Acknowledgments

We thank Kenway Louie, Tony Movshon, Alex Williams, and members of the Constantinople lab for feedback on the manuscript and helpful discussions.

## Author Contributions

S.S.S. collected electrophysiology data, with assistance from M.L.D. and R.M.W. S.S.S. performed muscimol experiments, and analyzed electrophysiology and behavioral data. D.P. developed the hLDS, under the supervision of D.H. and C.S. D.H. performed mutual information analysis. A.M. contributed to behavioral modeling. S.S.S, D.P. and C.M.C. prepared the figures. C.M.C. and S.S.S wrote the manuscript. C.M.C. and C.S. supervised the project.

## Data Availability

The data generated in this study will be deposited in a Zenodo database upon publication.

## Code Availability

Code used to analyze all data and generate figures will be available at https://github.com/constantinoplelab/published/tree/main upon publication.

## Supplemental materials

**Figure S1:**
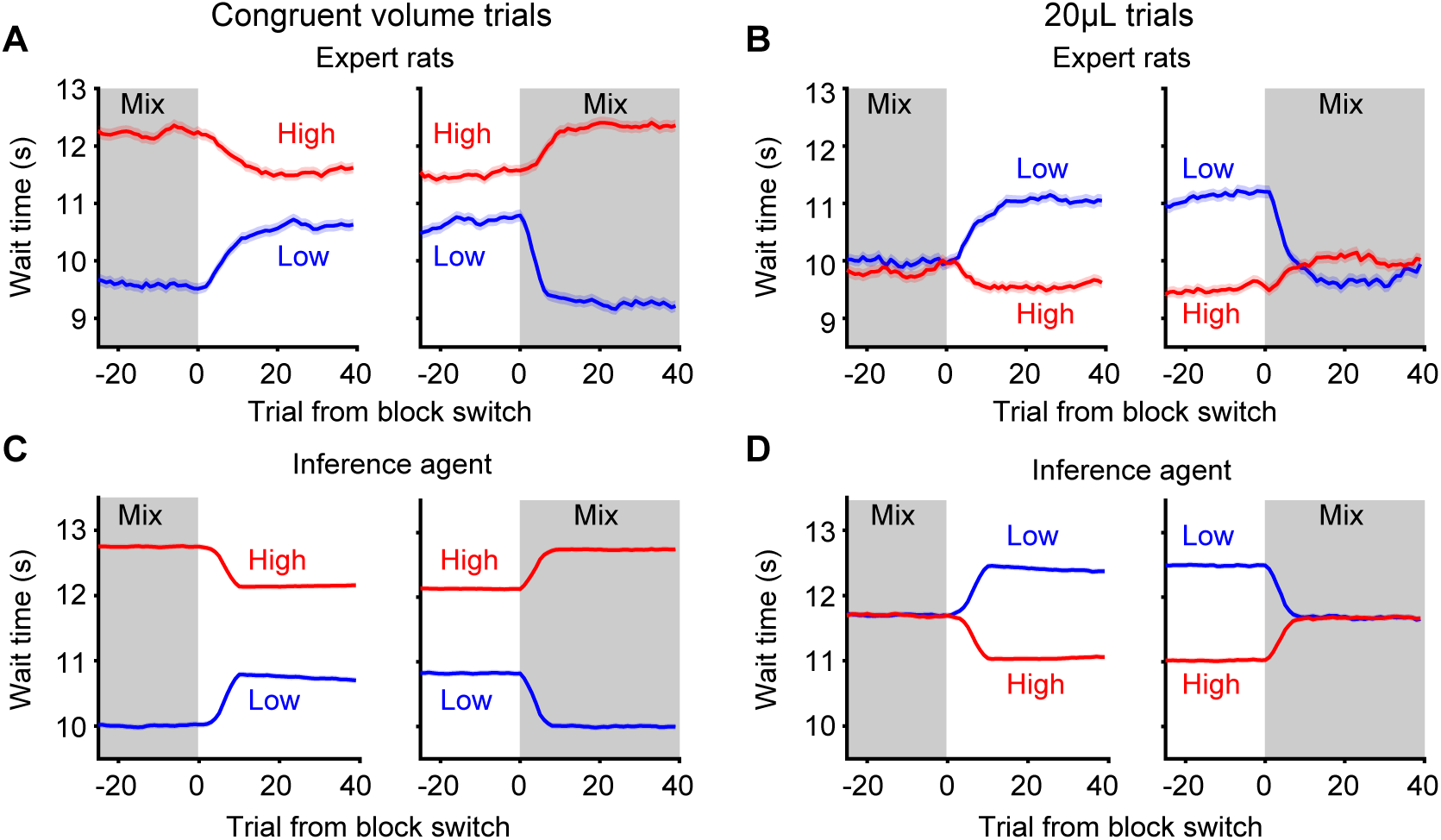
Wait time transition dynamics, related to Figure 1. **A.** Mean wait time dynamics for congruent volume trials. Mixed to high and high to mixed transitions include 40 and 80 *µ*L trials. Mixed to low and low to mixed transitions include 5 and 10 *µ*L trials. Wait times are influenced by both block and volume. Expert rats wait longer for large volume rewards than small volume rewards in mixed blocks. Wait times for the same volumes decrease when rats transition to a block with a higher opportunity cost (mixed to high and low to mixed). Wait times for the same volumes increase when rats transition to a block with a lower opportunity cost (mixed to low and high to mixed). **B.** Mean wait time dynamics for 20*µ*L trials. Wait times increase when the opportunity cost decreases. Wait times decrease when the opportunity cost increases. **C-D.** Block inference model recapitulates wait time changes depending on both block and volume.

**Figure S2:**
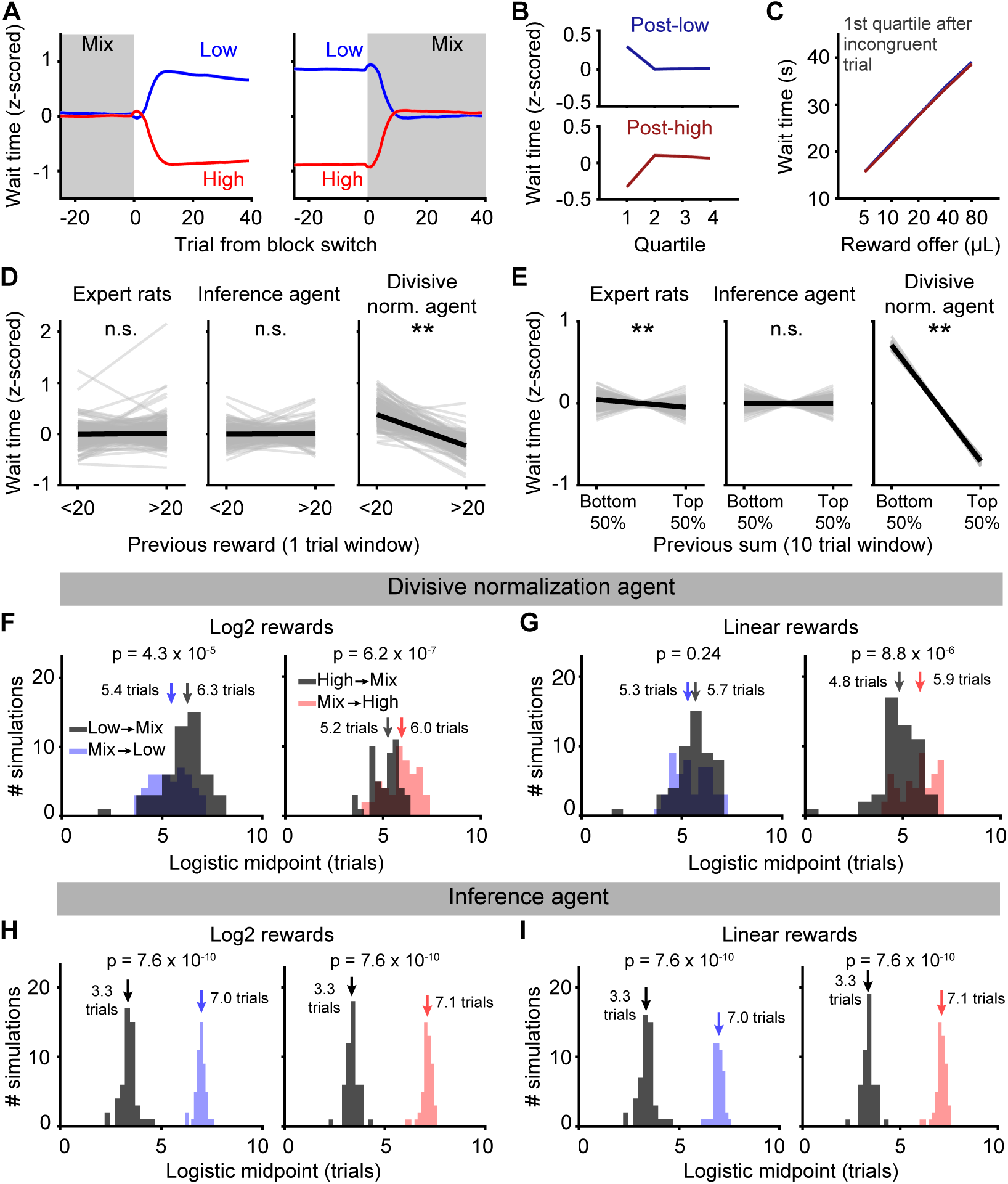
Divisive normalization agent with shorter integration windows, related to Figure 1. **A.** We simulated the behavior of a divisive normalization agent that integrated over 10 trials. The model was simulated for the trial sequences of each rat, and then predicted wait times were averaged over simulations (i.e., n=349 simulated agents). Data are mean +/- s.e.m. **B.** The divisive normalization agent also predicts rapid changes in wait times at blocks transitions, within the first quartile. **C.** With a short integration window, the divisive normalization agent can also adjust to the mixed block rapidly after the first incongruent trial. **D.** Mean z-scored wait times for 20 *µL* offers in mixed blocks conditioned on whether the previous reward was greater or less than 20 *µL* for expert rats (*p* = 0.69), an inference agent (*p* = 0.39), and a divisive normalization agent (*p* << 0.001). The divisive normalization model, but not inference model, predicts modulation of wait times based on the previous reward. All p-values from Wilcoxon signed-rank test. **E.** Given that divisive normalization integrates over multiple trials, we also computed the sum of the ten rewards preceding each 20*µ*L trial in a mixed block. Wait times for 20*µ*L in mixed blocks following sequences of large or small rewards (top and bottom 50% of the distribution) showed significant but small differences that could reflect mistaken inferences following sequences of small or large rewards for expert rats (*p* << 0.001). The divisive normalization model (*p* << 0.001), but not inference model (*p* = 0.54), predicts strong modulation of wait times based on the sum of the previous 10 rewards. All p-values from Wilcoxon signed-rank test. **F.** We fit logistic functions to block transitions of 50 simulated divisive normalization agents. In contrast to what was observed in rats, the model predicted faster behavioral changes for mixed-into-low transitions (5.4 trials) compared to low-into-mixed (6.3 trials), when rewards were represented on a log2 scale (which was used throughout the paper). The model predicted faster changes for high-to-mixed transitions (5.2 trials) compared to mixed-to-high transitions (6.0 trials, *p* = 6.2 × 10^−7^). All p-values from Wilcoxon signed-rank test. **G.** Same as panel F, but the model simulated wait times using rewards on a linear scale. In this case, there was no significant difference between low-into-mixed (5.7 trials) and mixed-into-low transitions (5.3 trials, *p* = 0.24), which was inconsistent with rats’ behavior. Transitions into mixed blocks from high blocks (4.8 trials) were still significantly faster than mixed blocks into high blocks (5.9 trials, *p* = 8.8 × 10^−^6). All p-values from Wilcoxon signed-rank test. This model therefore predicts faster transitions into low blocks, likely due to the nonlinearity in the divisive normalization computation. **H.** Similar to expert rats, the inference model predicts that transitions into mixed blocks, which contain incongruent trials, should be faster from both low (3.3 vs. 7.0 trials; *p* = 7.6 × 10^−10^) and high blocks (3.3 vs 7.1 trials; *p* = 7.6 × 10^−10^). All p-values from Wilcoxon signed-rank test. **I.** The inference model made identical predictions for rewards on a linear scale.

**Figure S3:**
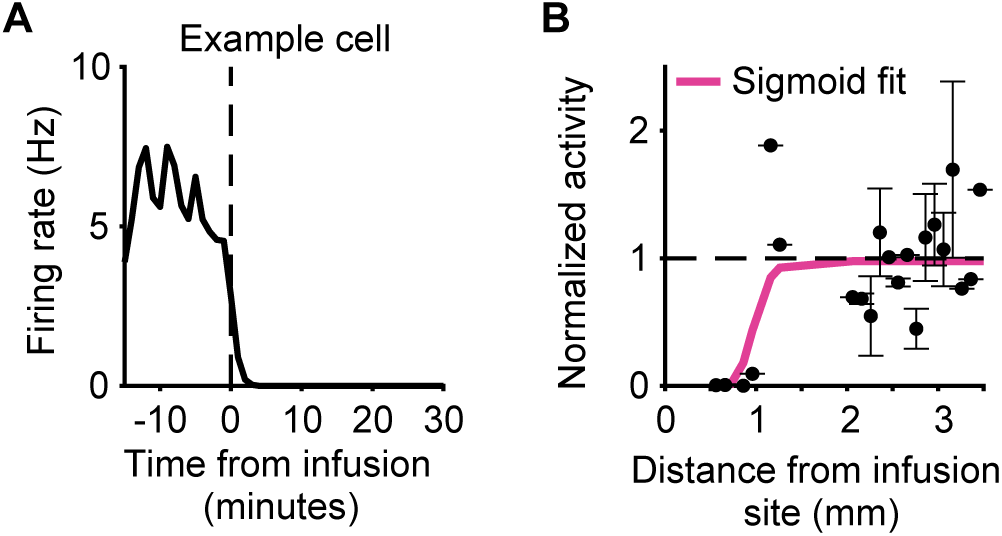
Muscimol inactivation of lateral OFC, related to Figure 2. **A.** Example neuron recorded within the infusion radius. The neuron is completely silenced within minutes after the infusion. **B.** Average firing rate for neurons in 0.1 mm bins after muscimol infusion. Post-infusion firing rates were normalized to pre-infusion firing rates. We fit a sigmoid to the data, and the midpoint was 0.8mm. Error bars are standard deviation.

**Figure S4:**
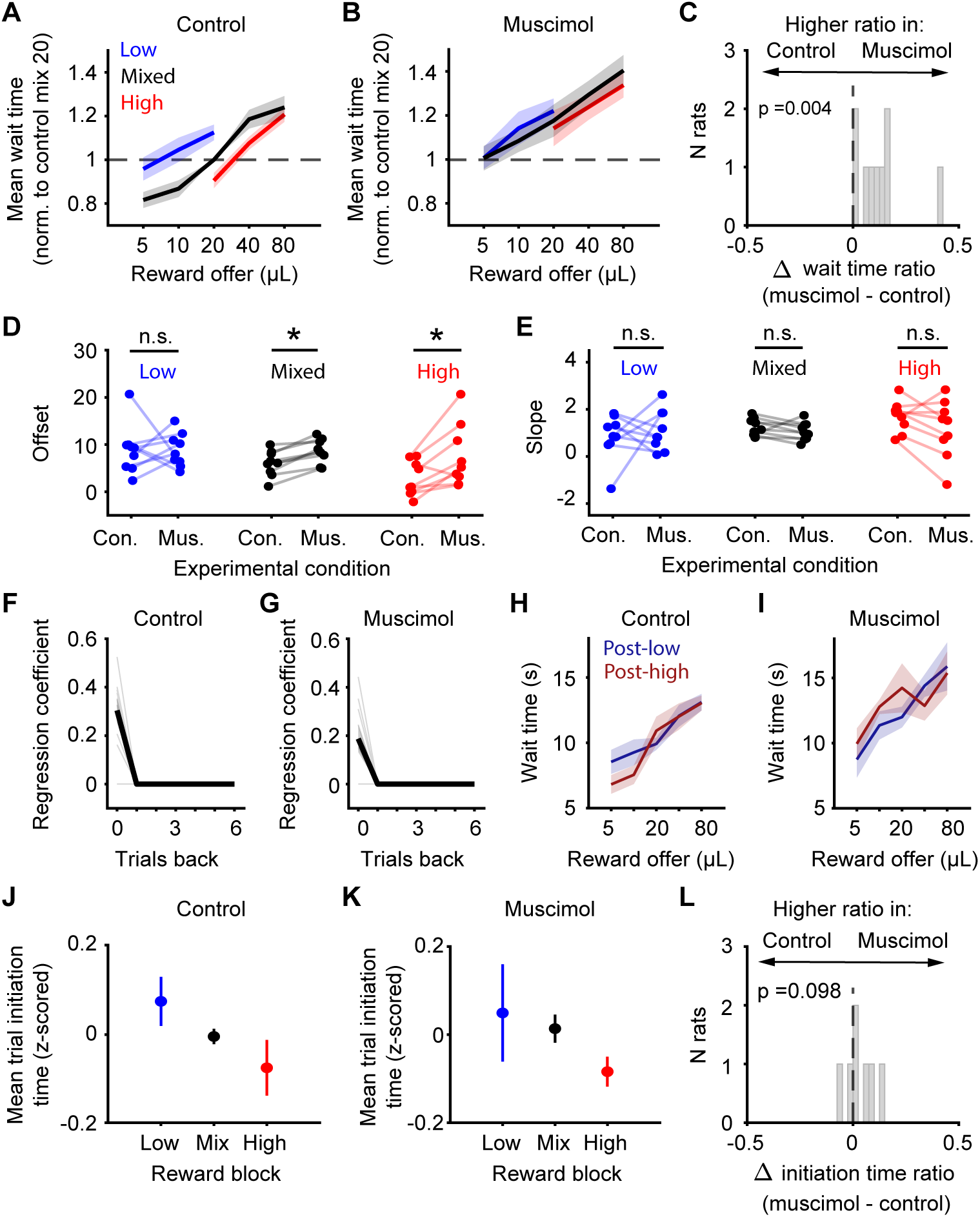
OFC inactivation impairs wait time sensitivity to hidden reward states but not volumes or previous rewards, related to Figure 2. **A,B.** Mean (+/- s.e.m) wait times normalized to 20µL mixed block trials in control sessions (**A**) and muscimol sessions (**B**). **C.** Difference in wait time ratios (high 20µL/low 20µL) between muscimol and control sessions for each rat. A higher wait time ratio (closer to 1) indicates that wait times on 20µL trials were more similar between blocks. *p* = 0.004, Wilcoxon signed-rank test. **D,E.** Offset parameter (**D**) and slope parameter (**E**) from linear regression of wait time against reward volume for each block. Each dot is 1 rat. Offset p-values: low *p* = 0.91, mix *p* = 0.004, high *p* = 0.02. Slope p-values: low *p* = 0.91, mix *p* = 0.13, high *p* = 0.16. Wilcoxon signed-rank test. **F,G.** Regression coefficients for wait time in control and muscimol sessions. Wait times were regressed against reward offers on the previous trial in mixed blocks only. **H,I.** Wait times in the first quartile of mixed blocks after the first incongruent trial, conditioned on the previous block type. Wilcoxon signed-rank text, Control: 5µL *p* = 0.63, 10µL *p* = 0.63, 20µL *p* = 0.38, 40µL *p* = 0.94, 80µL *p* = 0.69; Muscimol: 5µL *p* = 1, 10µL *p* = 0.38, 20µL *p* = 0.63, 40µL *p* = 1, 80µL *p* = 1. **J-K**. Mean (+/- s.t.d) z-scored trial initiation times for each block (averaged over all volumes) in control (**J**) and muscimol (**K**) sessions. **L**. Difference in trial initiation time ratios (high /low) between muscimol and control sessions for each rat. Trial initiation time ratios were not significantly different between muscimol and control sessions (*p* = 0.098, Wilcoxon signed-rank test.)

**Figure S5:**
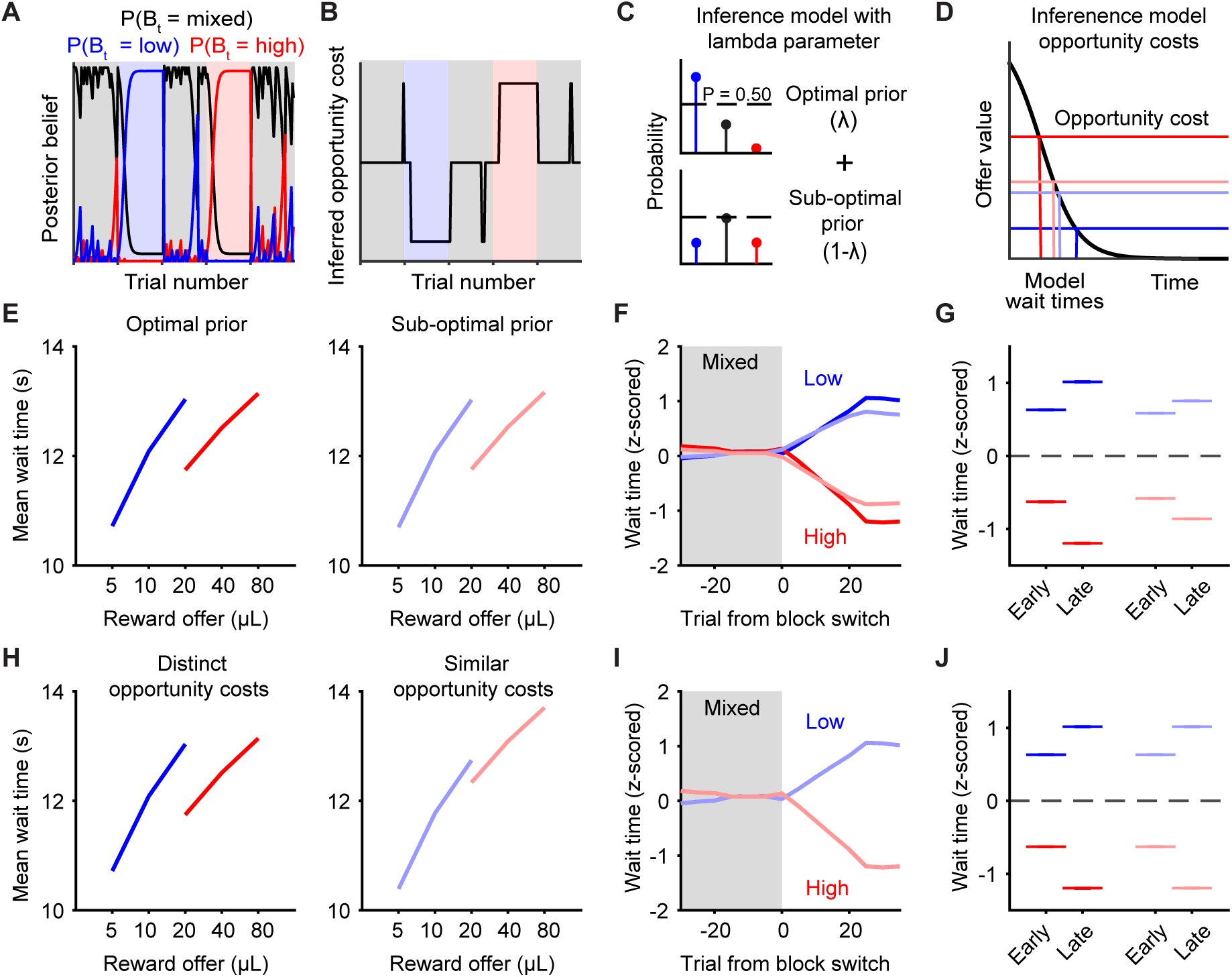
Other models are unable to capture OFC inactivation effects, related to Figure 2. **A.** Inference model computes a posterior probability for each block on each trial. The model predicted block is the one with the highest posterior probability. **B.** Example model-inferred opportunity cost selected based on the maximum posterior belief in **A**. The model selects one of three distinct opportunity costs, depending on the most likely block. Offer values on each trial are compared to that block-specific opportunity cost. **C.** Inference model with lambda parameter schematic. The model generates the sub-optimal prior by weighting between the true, optimal prior, and an uninformative prior. The optimal prior is based on knowledge of recent rewards, block transition probabilities, etc. The uninformative prior only incorporates knowledge of the overall block probabilities (0.5 for mixed blocks, 0.25 for low and high blocks). **D.** Schematic showing how changes in the opportunity cost parameter for each block change model predicted wait times. **E.** Simulated wait times with the inferential model using an optimal prior and a sub-optimal prior. A sub-optimal prior results in mistaken inferences, but to a similar degree in both high and low blocks, and therefore does not strongly reduce the wait time ratio. **F,G.** Simulated mean z-scored wait times early (trials 15-20) and late (trials 35-40) in a block after transitions from a mixed block. Dark colors indicate an optimal prior, light colors indicate a sub-optimal prior. A sub-optimal prior does not change the transition dynamics (i.e., midpoint of the sigmoid), but weakly changes the asymptote late in the block. **H.** Simulated wait times using the inferential model with a distinct opportunity cost associated with each block or similar opportunity costs associated with each block. An agent with more similar opportunity costs exhibits weaker contextual effects. **I,J.** Simulated mean changes in z-scored wait times early (trials 15-20) and late (trials 35-40) in a block after transitions from a mixed block. Dark colors indicate distinct opportunity costs, light colors indicate similar opportunity costs. Using similar opportunity costs for each block does not change the transition dynamics. Asterisks indicate significant differences between mean z-scored wait times for low and high blocks (*p* << 0.001 for all comparisons, Wilcoxon sign-rank test).

**Figure S6:**
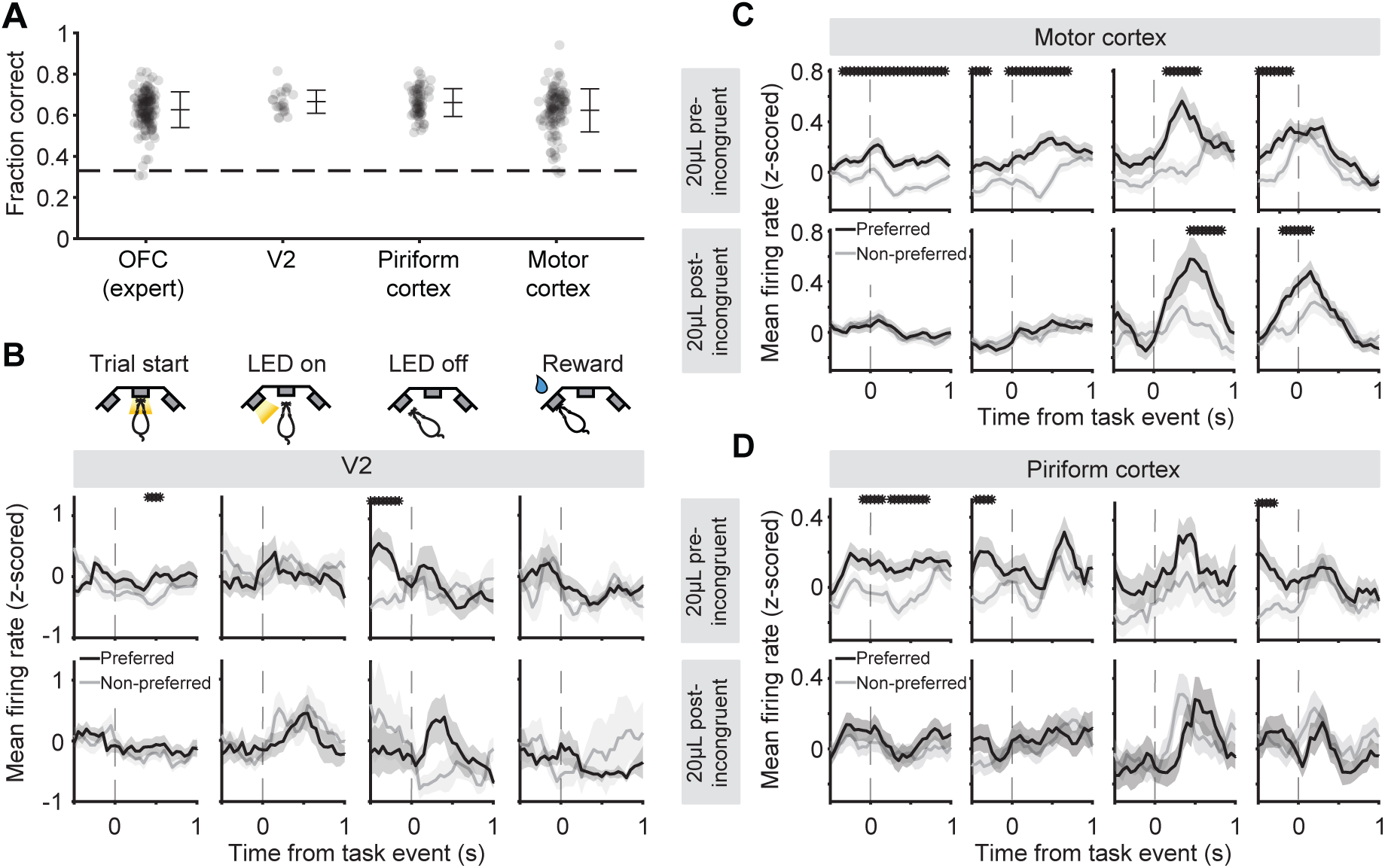
Responses to incongruent trials are not ubiquitous, related to Figure 3. **A.** Multinomial logistic regression decoder performance for each region. Each dot is one recording session. Error bar is mean +/- s.t.d. **B.** Mean (+/- s.e.m) firing rates for motor cortex (M1 or M2) neurons with significant block sensitivity on 20*µL* trials preceding (*top*) versus following (*bottom*) the first incongruent trial in a mixed block (Trial start: 1123/2907 neurons, LED on: 762/2907 neurons, LED off: 452/2907 neurons, Reward: 789/2907). Asterisks indicate a p-value *<* 0.05, non-parametric permutation test. **C.** Same as A but for piriform cortex (Trial start: 438/1522 neurons, LED on: 313/1522 neurons, LED off: 199/1522 neurons, Reward: 285/1522 neurons). Asterisks indicate a p-value *<* 0.05, non-parametric permutation test. **D.** Same as A but for V2 (Trial start: 117/266 neurons, LED on: 56/266 neurons, LED off: 39/266 neurons, Reward: 45/266 neurons). Asterisks indicate a p-value *<* 0.05, non-parametric permutation test.

**Figure S7:**
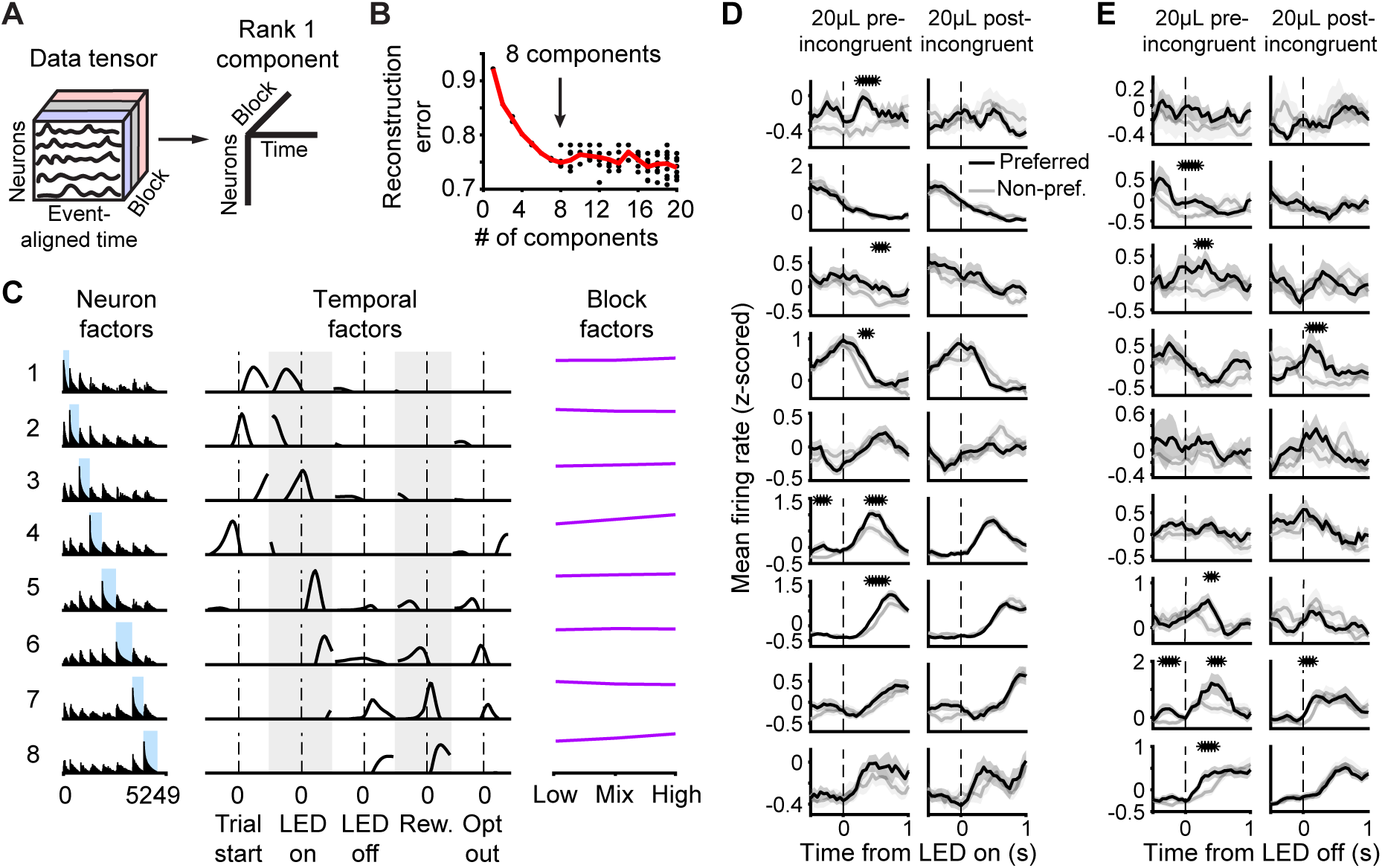
Tensor components analysis, related to Figure 4. **A.** TCA schematic. **B.** To determine the dimensionality, or rank, that should be applied to the neurophysiology data, we iteratively tried different numbers of dimensions, or ‘tensor components’, and computed the model reconstruction error. More than 8 components failed to improve model performance (elbow method). **C.** Neuron factors, temporal factors, and block factors for the TCA model of rank 8 for expert rats in OFC. Highlighted regions indicate neurons with a maximum loading for the given cluster. **D.** Mean (+/- s.e.m) z-scored firing rates for neurons in each cluster with significant block sensitivity on 20*µ*L trials preceding (*left*) versus following (*right*) the first incongruent trial in a mixed block, aligned to LED on. Asterisks indicate a p-value *<* 0.05, non-parametric permutation test. **E.** Same as panel D but aligned to LED off.

**Figure S8:**
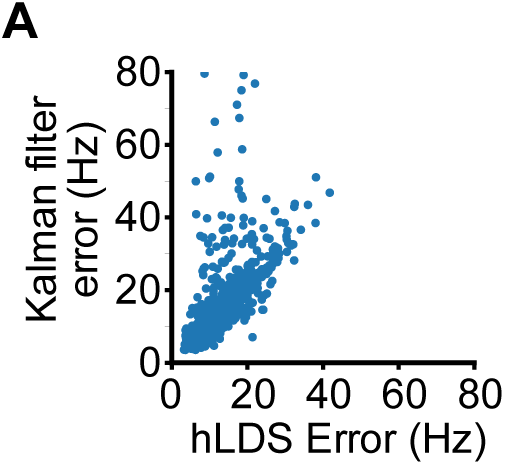
Hierarchical LDS outperforms dimensionality-matched Kalman filter, related to Figure 5. **A.** Given that the hLDS was fit with an 11 dimensional latent space (10 fast latents, 1 slow latent), an 11-dimensional standard Kalman Filter was also fit to each session. The reconstruction error for left-out neurons on held-out test data was compared across models in recordings from expert rats, and favored the hLDS. *p* << 0.001, Wilcoxon sign-rank test.

**Figure S9:**
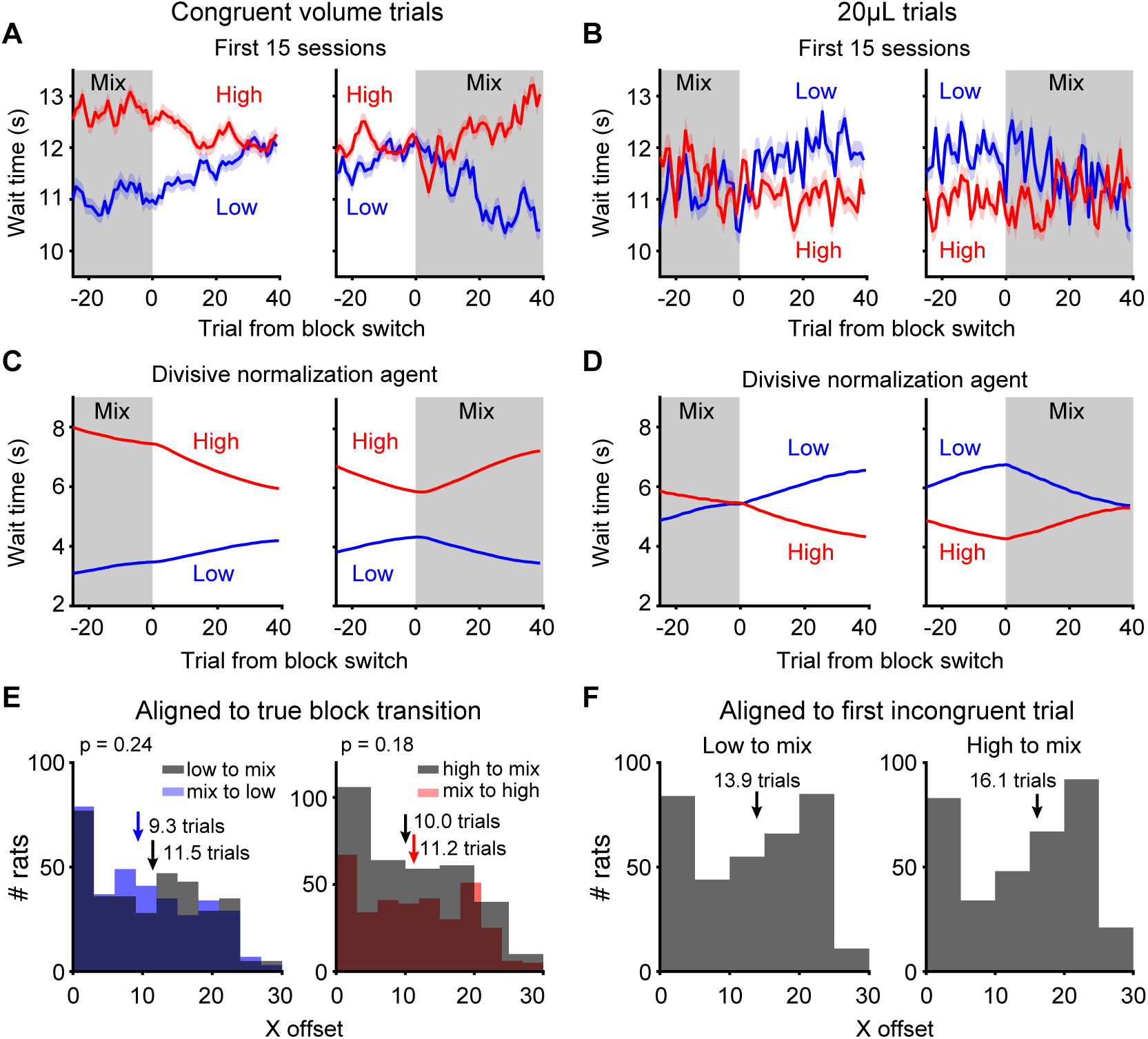
Raw wait time dynamics early in training are captured by the divisive normalization model, related to Figure 6. **A.** Mean wait time dynamics for congruent volume trials. Mixed to high and high to mixed transitions include 40 and 80 *µ*L trials. Mixed to low and low to mixed transitions include 5 and 10 *µ*L trials. Wait times are influenced by previous rewards, changing gradually over time. **B.** Mean wait time dynamics for 20*µ*L trials. **C-D.** Divisive normalization model recapitulates wait time changes dependent on previous rewards for congruent volume trials and 20*µ*L trials. **E.** The distribution of x values (number of trials after a block transition) that corresponded to the midpoint of each logistic fit, for different block transitions. Arrows show median values, p values are for Wilcoxon signed-rank test comparing the two distributions in each plot. Statistically different x-offset values indicate that rats were faster to adjust their wait times at transitions into mixed blocks, compared to transitions into high or low blocks. **F.** Distribution of x-offset values as in panel E, but when z-scored wait times were aligned to the first incongruent trial in mixed blocks. Arrows indicate median values of each distribution. Incongruent trials do not exist at transitions into high or low blocks, which is why those block transitions are not included in this plot.

**Figure S10:**
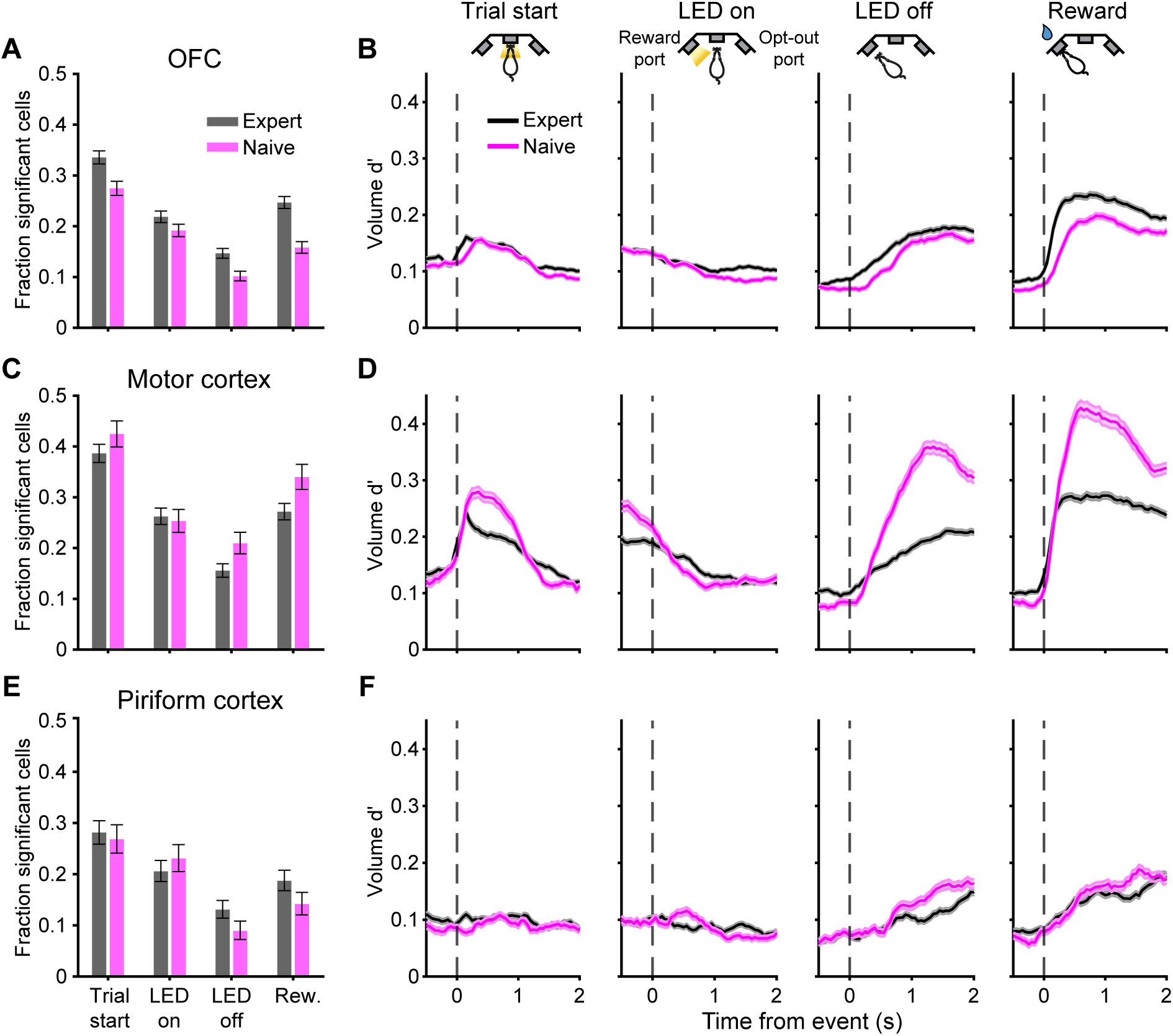
Encoding of reward volumes, related to Figure 7. **A.** Fraction of OFC cells with significant encoding of low versus high blocks (using all volumes) at each task alignment for both expert and naive rats. These cells were included in the analyses comparing pre-incongruent and post-incongruent firing rates on 20*µ*L trials. Naive data are combined over all recording sessions. Error bars are binomial confidence intervals. **B.** Mean encoding of reward volume (discriminability or d’; 5/10*µ*L versus 40/80*µ*L rewards) for OFC cells in mixed blocks for expert rats (black) and block naive rats (pink). **C.** Same as panel A but for motor cortex cells. **D.** Same as panel B but for motor cortex cells. **E.** Same as panel A but for piriform cortex cells. **F.** Same as panel B but for piriform cortex cells.

